# Structure of a dopamine-binding RNA aptamer reveals metal-mediated ligand recognition

**DOI:** 10.64898/2026.06.02.729643

**Authors:** Shea H. Siwik, David Stelzig, Simone D. Hall, Robert T. Batey

**Affiliations:** Department of Biochemistry, University of Colorado, Boulder, CO 80309-0596, USA

## Abstract

The selection of small-molecule binding RNA aptamers enables the creation of ligand-responsive RNA tools, yet most aptamers fail to operate reliably across diverse environments. Scaffolded selection addresses this limitation by preserving the tertiary architecture of a riboswitch scaffold while driving the evolution of a new ligand-binding pocket. Using the *xpt* purine riboswitch aptamer, we previously generated dopamine-binding aptamers. Here, we report the crystal structures of two representative variants in their apo and dopamine-bound states to define how they recognize ligand while maintaining scaffold integrity. The structures demonstrate that scaffolded selection enforces global fold conservation and retains the defining tertiary interactions of the parental riboswitch. Local remodeling, triggered by deletions acquired during selection, rewires the three-way junction to build extensive interaction networks that host the dopamine-binding pocket. A deeply buried potassium ion anchors ligand recognition by coordinating the dopamine hydroxyl group while the RNA engages the catechol ring through stacking and hydrogen bonding interactions. Structure probing shows minimal conformational changes upon ligand binding, indicating that the aptamers adopt a largely preorganized fold. These findings strengthen the central premise of scaffolded selection: riboswitch-derived “superfolder” architectures can bias *in vitro* selections towards aptamers that conserve global structure while supporting locally diverse binding pockets. This balance between structural stability and local plasticity expands the capacity of a single RNA fold to recognize chemically distinct ligands and positions scaffolded selection as a powerful platform for engineering robust RNA-based sensing and regulatory devices.

**TOC Abstract (graphical):** 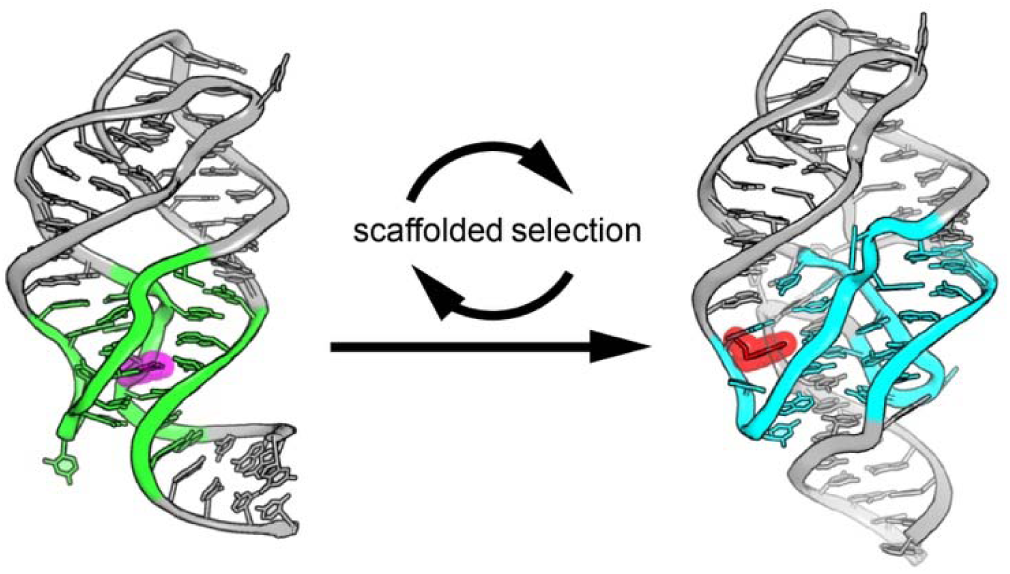

## INTRODUCTION

Small molecule-binding nucleic acid aptamers hold substantial promise for a wide range of biotechnological and medical applications (1–4). These aptamers can be efficiently generated using well-established *in vitro* selection strategies (5–8) and readily incorporated into functional devices whose activity is regulated by small molecule binding. Considerable effort has focused on developing small molecule-responsive systems that control gene expression at the mRNA level. In one widely explored strategy, an aptamer is embedded within an alternative splicing cassette placed within a mammalian mRNA (9–12). In the absence of ligand binding, full exon inclusion introduces a premature stop codon that triggers nonsense mediated decay, whereas ligand binding promotes an alternative splicing pattern that permits expression of a therapeutic protein. Beyond gene regulation, small molecule-binding aptamers are increasingly used as key components of biosensors, including fluorescent RNA sensors that detect metabolites in living cells and generate a fluorescent readout (1,13).

Despite the development efforts spanning more than three decades, only a small fraction of RNA aptamers have been successfully translated into robust, widely used platforms. The theophylline aptamer remains a notable exception, exhibiting reliable performance across diverse platforms and consequently achieving widespread adoption (14). Together, these observations highlight a critical unmet need for strategies that enable the systematic development of small molecule-binding aptamers with robust function across varied cellular and technological contexts.

In contrast to aptamers selected *in vitro*, bacteria have evolved a diverse collection of highly functional natural aptamers. These aptamers are most commonly found within riboswitches, regulatory RNA elements typically located in the 5’ leader regions of bacterial mRNAs that control downstream expression by selectively binding a cognate metabolite through a structured aptamer domain (15–18). To date, more than 55 classes of riboswitches have been experimentally validated, collectively recognizing a wide range of ligands, including nucleobases, amino acids, cofactors, second messenger molecules, and metal ions (19). Underscoring their robustness, riboswitches are abundant across bacterial phyla with Firmicutes and Fusobacteria relying extensively on riboswitch-mediated regulation of essential housekeeping genes (20,21). Their widespread success reflects strong evolutionary selection for aptamer domains that fold rapidly and reproducibly: properties that have led riboswitch aptamers to be described as “superfolders” (22). This rapid and faithful folding is critical because many riboswitches function co-transcriptionally. While RNA polymerase remains in the 5’-leader sequence, the nascent RNA must quickly fold, bind its ligand, and adopt the appropriate regulatory conformation to determine the fate of the transcript (15,22,23).

Because *in vitro* selection directly enriches only for ligand binding, the robust folding of newly isolated aptamers is not guaranteed, frequently limiting their functional deployment. Modified selection strategies seek to overcome this limitation by drawing inspiration from the intrinsically stable architectures of riboswitch aptamer domains. In one such approach, a well-folding RNA scaffold is used to promote efficient folding of the selected aptamer (24). The most widely used scaffold is derived from the purine riboswitch aptamer domain that adopts a common tertiary architecture in biological RNAs: a three-way junction (3WJ) stabilized by a distal tertiary interaction (25,26). Functional RNAs that use this architectural theme include the hammerhead ribozyme (27,28), multiple riboswitch aptamer domains (29–31), and folding and assembly units of ribonucleoprotein particles (32,33).

In this scaffolded selection strategy, the binding pocket embedded within the 3WJ of the parent RNA is fully randomized, while the surrounding secondary and tertiary architecture is preserved (**Figure 1**) (24). This approach seeks to repurpose the 3WJ to bind new ligands while maintaining the global fold and elements that confer robust folding. Using this strategy, the *Bacillus subtilis xpt-pbuX* riboswitch has been evolved to bind to 5-hydroxytryptophan (5HTP)/serotonin (24), L-3,4 hydroxyphenylalanine (L-DOPA)/dopamine (24), quinine (34), caffeine (34), and theophylline (35). A related strategy employing the *Vibrio vulnificus add* adenine riboswitch enabled the recognition of a fluorogenic small molecule (36,37). Available structural data reveals preservation of the parent aptamer fold, underscoring the effectiveness of this approach in achieving productive RNA folding (24,37,38). Collectively, these studies demonstrate the utility of scaffolded selection for generating novel small molecule-binding aptamers.

**Figure 1.**
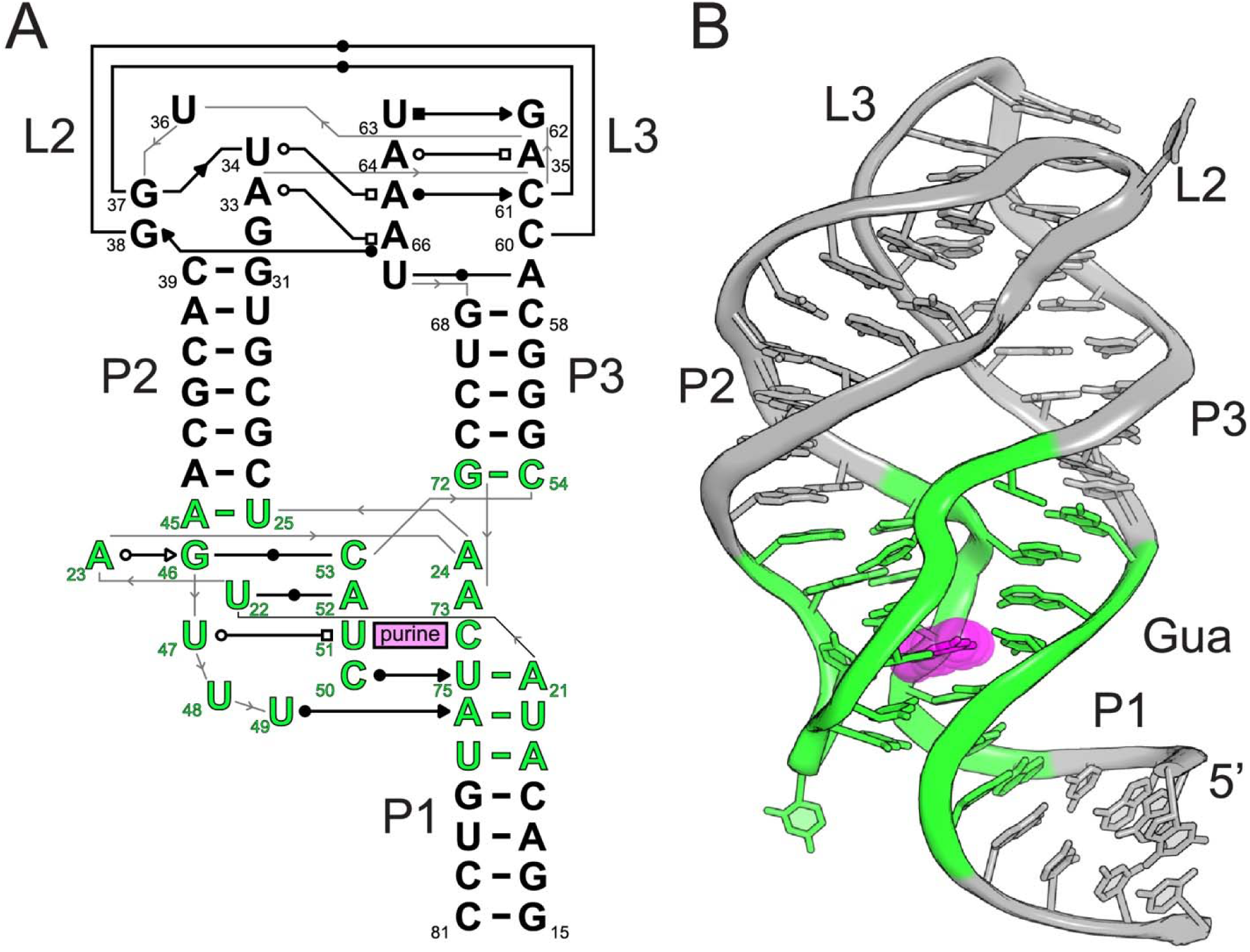
Parental *B. subtilis xpt* guanine riboswitch. (A) Secondary structure of the *B. subtilis xpt* guanine riboswitch. Base-mediated interactions denoted using the standard Leontis-Westhof notation (84), and the numbering is consistent with that used in the crystallographic models of the *B. subtilis xpt* aptamer (65). Nucleotides whose identities were fully randomized in the initial L-DOPA selection are highlighted in green. (B) Tertiary structure of the *xpt* riboswitch bound to guanine (magenta) (85). Nucleotides within an 8 Å distance of guanine were chosen for randomization (green) (24).

To further elucidate how scaffolded selection repurposes riboswitch architectures to generate new functional aptamers, we determined the structures of two members of the class I L-DOPA/dopamine (DGR-1) aptamer family (24). These structures show that the parental 3WJ architecture is preserved and that ligand binding occurs within a pocket created by one of the joining strands, accommodating a single molecule of dopamine. Although the global 3WJ is conserved, the P1 and P3 helices have been altered through sequence rearrangement during the selection. The DGR-1 aptamer primarily recognizes the catechol moiety of dopamine with ligand engagement mediated by a coordinating cation that bridges the RNA and small molecule. Comparisons of the apo and ligand-bound structures indicate that the RNA is largely pre-organized for ligand recognition which is supported by selective 2’-hydroxyl acylation analyzed by primer extension (SHAPE) chemical probing of both aptamers. Together, these results expand upon the growing family of natural and synthetic purine family aptamers and demonstrate how a conserved RNA fold can be repurposed to selectively recognize chemically distinct small molecule ligands.

## METHODS

### RNA synthesis and purification

DNA oligonucleotides were synthesized by Integrated DNA Technologies (IDT) and amplified by recursive polymerase chain reaction (PCR) (39). Sequences of all DNA templates and RNA constructs used in this study are provided in **Supplementary Tables S1 and S2**. The 3’ DNA primer for template amplification of crystallization constructs contained two 2’-*O*-methoxy modifications at its 5’ end to promote RNA homogeneity and prevent non-templated nucleotide addition at the 3’ end of the transcript by T7 RNA polymerase (40,41).

dsDNA templates were transcribed *in vitro* using T7 RNA polymerase as described previously (42). Following transcription, reactions were buffer exchanged into Milli-Q H_2_O using a 10 kDa molecular weight cut off (MWCO) centrifugal concentrator (Amicon) and purified by 8% denaturing polyacrylamide gel electrophoresis (PAGE; 29:1 acrylamide:bis-acrylamide, 8 M urea, 1x TBE (90 mM Tris base, 90 mM boric acid, 3 mM ethylenediaminetetraacetic acid (EDTA))) (43). Product RNA bands were visualized by UV shadowing, excised and passively eluted into either 0.5x TE buffer (5 mM Tris, pH 8.0, 0.5 mM EDTA) or Milli-Q H_2_O. Eluted RNA was buffer exchanged into 0.5x TE buffer, concentrated using a 10 kDa MWCO centrifugal concentrator (Amicon), and quantified by absorbance at 260 nm. Extinction coefficients were calculated by using the IDT OligoAnalyzer tool (https://www.idtdna.com/calc/analyzer).

### Crystallization and structure determination

Structural studies were performed using the DGR-1A and DGR-1B L-DOPA/dopamine aptamer RNAs. DGR-1B RNA was crystallized at 20 °C by hanging drop vapor diffusion (44). Crystallization drops consisted of 1 µl of RNA-ligand solution (200 µM RNA and 400 µM dopamine or L-DOPA) mixed with 1 µl of precipitant solution containing 10% (v/v) 2-methyl-2,4-pentanediol (MPD), 20 mM spermine, 50 mM SrCl_2_, 50 mM LiCl, and 40 mM sodium cacodylate (pH 6.0) equilibrated over 500 µl of reservoir solution containing 35% (v/v) MPD. Crystals were cryoprotected by soaking in precipitant solution supplemented with 20% (v/v) MPD for 5 minutes at 20 °C and flash freezing in liquid nitrogen.

DGR-1A was co-crystallized with dopamine using hanging drop vapor diffusion by combining 1 µL of RNA solution (300 µM DGR-1A RNA and 1 mM dopamine hydrochloride) with 1 µL of the crystallization solution containing 11% (v/v) MPD, 8 mM hexammine cobalt(III) chloride, 12 mM NaCl, 80 mM KCl, and 40 mM MES ,pH 6.0, equilibrated over 500 µL of 35% (v/v) MPD. Crystals grew at 20 °C for 8 d prior to harvesting and flash freezing in liquid nitrogen.

X-ray diffraction data were collected at 100 K at the Macromolecular X-ray Crystallography Core at the University of Colorado Boulder with a RIGAKU XtaLAB MM003 and Dectris PILATUS 200K 2D hybrid pixel array detector (Cu Kα radiation, 1.54178 Å) and processed in HKL3000 (45). Additional datasets were collected at the Advanced Light Source (ALS) beamline 8.2.1 using a ADSC Quantum 315r detector (0.99992 Å) and processed with DIALS (46). Data collection and refinement statistics are reported in **Supplementary Table S3**.

The structure of the DRG-1B aptamer was solved by molecular replacement method using Phaser (47) and refined using PHENIX (48). Initial phasing of the DRG-1B structure used the three helices of the *xpt* guanine riboswitch (PDB ID: 4FE5) as search models (P1: nucleotides (nt) 15-21 and 75-81; P2 nt 25-31 and 39-45; P3 nt 54-59 and 67-72) (49). The remaining RNA was built into electron density using Coot (50) with iterative refinements of simulated annealing and energy minimization in PHENIX (48). The asymmetric unit contained two RNA molecules, which were built independently. RNA geometry was improved and validated using ERRASER (51) and MolProbity (52). The final DGR-1B model had R_work_ of 20.1% and R_free_ 24.9% with root-mean-square deviations (RMSD) of 0.002 Å for bond lengths and 0.58° for bond angles. Representative electron density of a composite omit 2F_o_-F_c_ map is shown in **Supplementary Figure S1A**.

For DGR-1A, the refined DGR-1B structure was used as the search model for molecular replacement in Phaser followed by rebuilding and refinement using Coot and PHENIX. The asymmetric unit contained two protomers of DGR-1A that were built independently with dopamine placed into the model manually. The final DGR-1A model had R_work_ of 23.3% and R_free_ 27.0% with RMSDs of 0.003 Å for bond lengths and 0.70° for bond angles. Representative electron density of a composite omit 2F_o_-F_c_ map is shown in **Supplementary Figures S1B** and **S2**.

### Isothermal Titration Calorimetry (ITC)

RNA for ITC experiments was transcribed *in vitro* using T7 RNA polymerase and purified under native conditions. Following transcription, reactions were treated TURBO DNase (0.5 U per µg of DNA template, Invitrogen) for 30 min at 37 °C followed by Proteinase K digestion (0.25 mg mL^−1^ final concentration, Invitrogen) for an additional 30 min at 37 °C (53). RNA was buffer-exchanged into 0.5x TE buffer using a 10 kDa MWCO centrifugal concentrator (Amicon) and dialyzed for 12 h at 4 °C against 500 mL of ITC buffer (50 mM 4-(2-hydroxyethyl)-1-piperazineethanesulfonic acid (HEPES)-KOH, pH 7.5, 50 mM KCl, 10 mM MgCl_2_, and 10 mM Tris(2-carboxyethyl)phosphine hydrochloride (TCEP-HCl)) or buffer-exchanged into ITC buffer three times using a 10 kDa MWCO centrifugal concentrator (Amicon). TCEP-HCl was used as dopamine rapidly oxidizes resulting in a product that does not interact with the RNA (54–56).

RNA concentrations were determined by absorbance at 260 nm using calculated molar extinction coefficients. Ligand concentrations were determined using published molar extinction coefficients where available. For ligands lacking reported values, extinction coefficients were determined experimentally (**Supplementary Table S4**).

ITC measurements were conducted using a MicroCal iTC200 (Malvern) or a Low Volume Affinity ITC (Waters TA) at 25 °C using established protocols (57). Experimental c-values were selected to enable accurate determination of dissociation constants (58). For data collected on the MicroCal iTC200, data were background-corrected by subtracting the average heat of ligand dilution into buffer and fit to a single-site model using Origin software. For low-affinity interactions, experiments were performed at high ligand-to-RNA ratios with the binding stoichiometry fixed at n = 1 to allow estimation of the dissociation constant (58); enthalpic or entropic values were not extracted under these conditions. For data collected on the Low Volume Affinity ITC, data were background-corrected by subtracting the heat of ligand dilution into buffer for each injection and fit to the independent model using NanoAnalyze software. Representative thermograms are shown in **Supplementary Figure S3**.

For experiments examining monovalent cation identity, HEPES acid, MgCl_2_, and TCEP-HCl were combined, and the pH of the solution adjusted to 7.5 using the appropriate cation hydroxide. The corresponding chloride salt was added to yield a final monovalent cation concentration of 125 mM cation (K^+^, Na^+^, Li^+^ or Cs^+^). Final buffer concentrations were 50 mM HEPES (pH 7.5), 10 mM MgCl_2_, 10 mM TCEP-HCl, and 125 mM (M^+^Cl^−^). RNA samples were dialyzed against three successive 500 mL buffer exchanges at 4 °C (8 hours per exchange) prior to ITC measurements. Complete thermodynamic values from ITC measurements of DGR aptamers binding dopamine, L-DOPA, and (*S*)-2-amino-3-(3,4-dihydroxyphenyl)propanamide (L-DOPA amide) are shown in **Supplementary Table S5**.

### Chemical footprinting analysis of RNA

DNA templates incorporating a 3′ SHAPE cassette were amplified by PCR followed by *in vitro* transcription and gel purification as previously described (59). RNA was purified by two rounds of 8% denaturing PAGE. After the final gel purification, the RNA was eluted into 0.5x TE at 4 °C for 2 h, buffer exchanged into 0.5x TE using a 10 kDa MWCO centrifugal concentrator, and stored at -70 °C.

Prior to SHAPE chemical probing, RNA was denatured at 95 °C for 3 min and snap-cooled on slushy ice for 5 min. Folding reactions used a 3x folding buffer containing 333 mM HEPES, pH 8.0, 20 mM MgCl_2_, and 333 mM K^+^, generated by adjusting the pH with the corresponding hydroxide. The folding buffers for the other monovalent cations were prepared in the same manner with the appropriate lithium, sodium, and cesium chloride salts and hydroxides. RNA, ligand, 3x folding buffer, and 0.5x TE were combined and incubated at 37 °C for 20 min.

1-methyl-7-nitroisatoic anhydride (1M7) was freshly prepared in anhydrous dimethyl sulfoxide (DMSO) and added to reactions that were incubated for an additional 20 min at 37 °C. Final reaction volumes (10 µL) contained 1x folding buffer, 0.2x TE, 1 pmol RNA, 1 mM dopamine, and 6.5 mM 1M7. Unless specified, reactions were conducted in buffer containing K^+^ as the monovalent cation. Control reactions contained neat DMSO instead of 1M7. Modified RNAs were stored at -20 °C prior to reverse transcription using a ^32^P-labeled primer without ethanol precipitation (60).

Reverse transcription products were resolved by 12% denaturing sequencing PAGE at 55 W. Gels were exposed to phosphor storage screens, imaged using a Typhoon (Amersham), and images quantified using the SAFA software package (61,62). Reactivities from DMSO controls were subtracted from 1M7-modified samples. Data were normalized by dividing each value by the average of the five highest reactivity values across all conditions. Each of three replicates was analyzed individually. Ligand-dependent changes in reactivity were calculated by subtracting -ligand values from +ligand values at each nucleotide position.

## RESULTS AND DISCUSSION

### Crystallization and structure determination of dopamine aptamers

To define the molecular basis for dopamine recognition by RNA, we solved atomic-resolution structures of a dopamine aptamer raised by scaffolded selection (24). We selected one of the dominant classes of dopamine aptamer generated by scaffolded selection (DGR-1) and two close related variants within the class (DGR-1A and DGR1-B) for crystallographic analysis. These aptamers share more than 90% sequence identity and are expected to adopt closely related structures (**Supplementary Figure S4**).

To facilitate crystallization, we modified the DGR aptamers in two ways. First, the length of the P1 helix was systematically varied to promote favorable lattice contacts which enabled crystallization of the *B. subtilis xpt-pbuX* guanine riboswitch (GR) in complex with hypoxanthine (63). Because the P2 and P3 helices of the GR scaffold participate in conserved tertiary interactions essential for proper folding (60), these helices were not altered. Second, an unpaired adenosine was appended to the 3′ end of the RNA to encourage helical stacking and intermolecular contacts within the crystal lattice.

To verify that these modifications did not impair ligand binding, we measured affinity for L-DOPA, the molecule used during the selection, using ITC. DGR-1A and DGR1-B bound L-DOPA with dissociation constants (K_D_) of 42 ± 4 µM and 150 ± 10 µM, respectively (**Table 1**). Given the close chemical similarity between L-DOPA and dopamine (chemical structures for all compounds used in this study are shown in **Supplementary Figure S5**), we also measured binding to dopamine and observed higher affinities for both aptamers (11 ± 1 µM for DGR-1A and 28 ± 1 µM for DGR-1B, **Table 1**). In both cases, the binding stoichiometry for L-DOPA/dopamine was consistent with a single ligand molecule binding per RNA. Further, the enthalpy/entropy compensation observed for the DGR aptamers is consistent with that observed for many natural riboswitches and synthetic aptamers (64) (**Supplementary Table S5**). These results confirmed that the crystallization-oriented modifications were not deleterious and supported use of both ligands for structural studies.

**Table 1:**
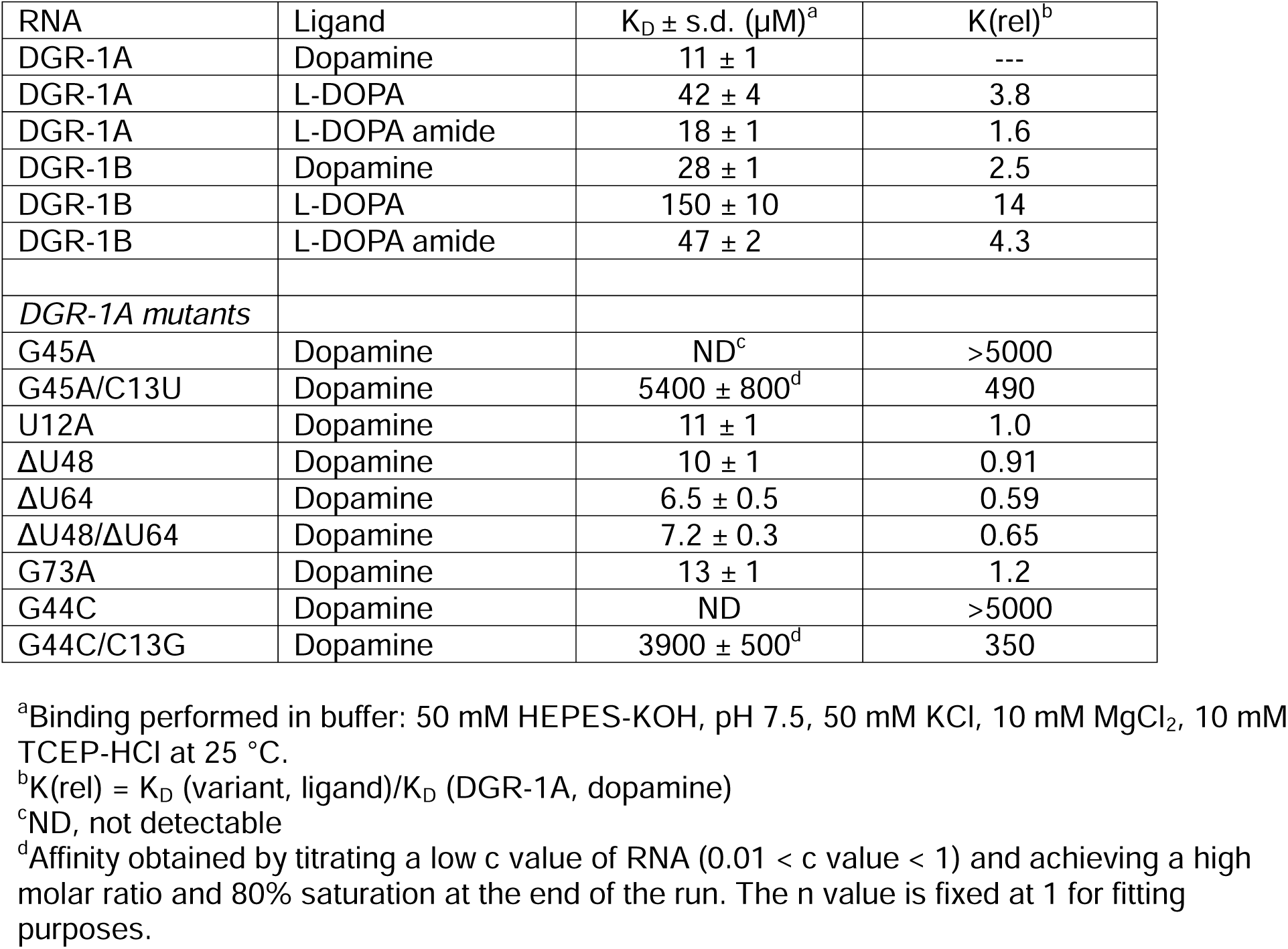
ITC analysis of DGR aptamers binding to ligands.

Diffraction-quality crystals were obtained for both aptamers in the presence of L-DOPA or dopamine. The DGR-1B RNA, containing a 6 base pair (bp) P1 helix, crystallized in the presence of L-DOPA and diffracted to 2.92 Å resolution. The structure was solved by molecular replacement with the three helices of the parental GR aptamer (PDB ID: 4FE5) and refined to a final R_work_/R_free_ of 0.201/0.249 (**Supplementary Table S3**). The asymmetric unit contained two protomers with nearly identical structures (RMSD = 1.46 Å). Unless otherwise stated, all structural descriptions refer to molecule A, which exhibited slightly improved geometry. Despite crystallization in the presence of L-DOPA, no electron density corresponding to the ligand was observed, indicating that DGR-1B represents the apo state.

DGR-1A, containing an 8 bp P1 helix, was co-crystallized in the presence of dopamine and diffracted to 2.59 Å resolution. The DGR-1B structure served as the molecular replacement model, yielding a final structure with an R_work_/R_free_ of 0.233/0.270. Two protomers were present in the asymmetric unit (RMSD = 0.740 Å), each containing clear electron density for a single dopamine molecule. Unless otherwise stated, all structural descriptions of DGR-1A refer to molecule A, representing the ligand-bound structure.

### The parental guanine riboswitch scaffold architecture is preserved in the DGR aptamers

A central premise of scaffolded selection is that the overall secondary and tertiary interactions of a highly evolved biological scaffold is maintained in the newly selected aptamers, promoting robust RNA folding (24). Members of the guanine/adenine riboswitch family share a conserved 3WJ architecture supported by a distal tertiary interaction between L2 and L3 (65–71). Both DGR-1A and DGR-1B aptamers preserve this global architecture with the P2 and P3 helices positioned side-by-side and P1 located beneath them (**Figure 2**).

**Figure 2.**
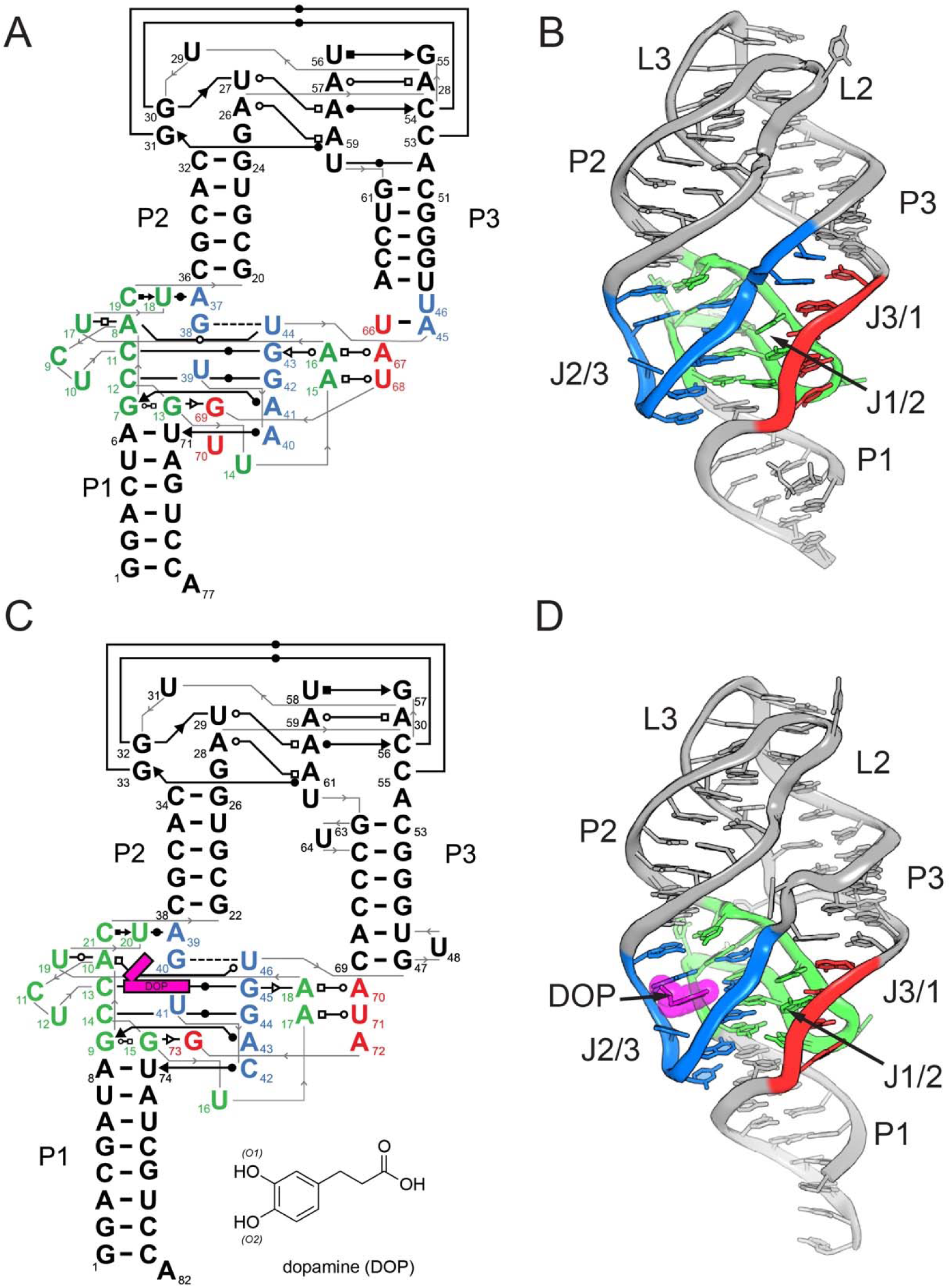
Architecture of the DGR-1A and DGR-1B aptamers. (A) Secondary structure of the DGR-1B aptamer with base-mediated interactions denoted using the standard Leontis-Westhof notation (84) and numbering consistent with that used in the crystal structure (12CJ PDB ID). The three joining strands are colored (green, J1/2; blue, J2/3; red, J3/1) to highlight the three-way junction region. (B) Cartoon representation of the three-dimensional structure of the DGR-1B aptamer using the same color scheme as in panel (A). (C) Secondary structure of the DGR-1A aptamer, with same notation scheme as in panel (A), but with numbering consistent with the crystal structure (12CI PDB ID). (D) Cartoon representation of the three-dimensional structure of the DGR-1A aptamer using the same color scheme as in panel (A) and dopamine (DOP) highlighted in magenta.

The principal difference relative to the parental GR scaffold lies in the orientation of P1. In the DGR aptamers, P1 is displaced from strict coaxial stacking with P3 and adopts an intermediate position between the P2 and P3 (**Figure 2B, D**). Expansion of the J1/2 sequence between P1 and P2 in the DGR aptamers likely disrupts the coaxial stacking found in the GR scaffold. Thus instead, in the DGR aptamers we observe continuous base stacking from P1 through the 3WJ into both P2 and P3. Despite this difference, the overall organization of the scaffold is maintained.

Global architecture is further enforced by conservation of the L2-L3 interaction. Nucleotides within the L2 and L3 loops are highly conserved in the guanine and adenine riboswitch classes and form extensive interactions that lock the two loops together (65,66). In the scaffolded selection of the DGR aptamers, L2, L3, and the paired regions adjacent to them were not randomized (24). In both DGR-1A and DGR-1B, these loops closely superimpose with their counterparts in the GR scaffold with a RMSDs of 0.36 Å and 0.73 Å, respectively (**Figure 3**). Together, these observations demonstrate that both global and local structural features defining the GR scaffold are retained following scaffolded selection.

**Figure 3.**
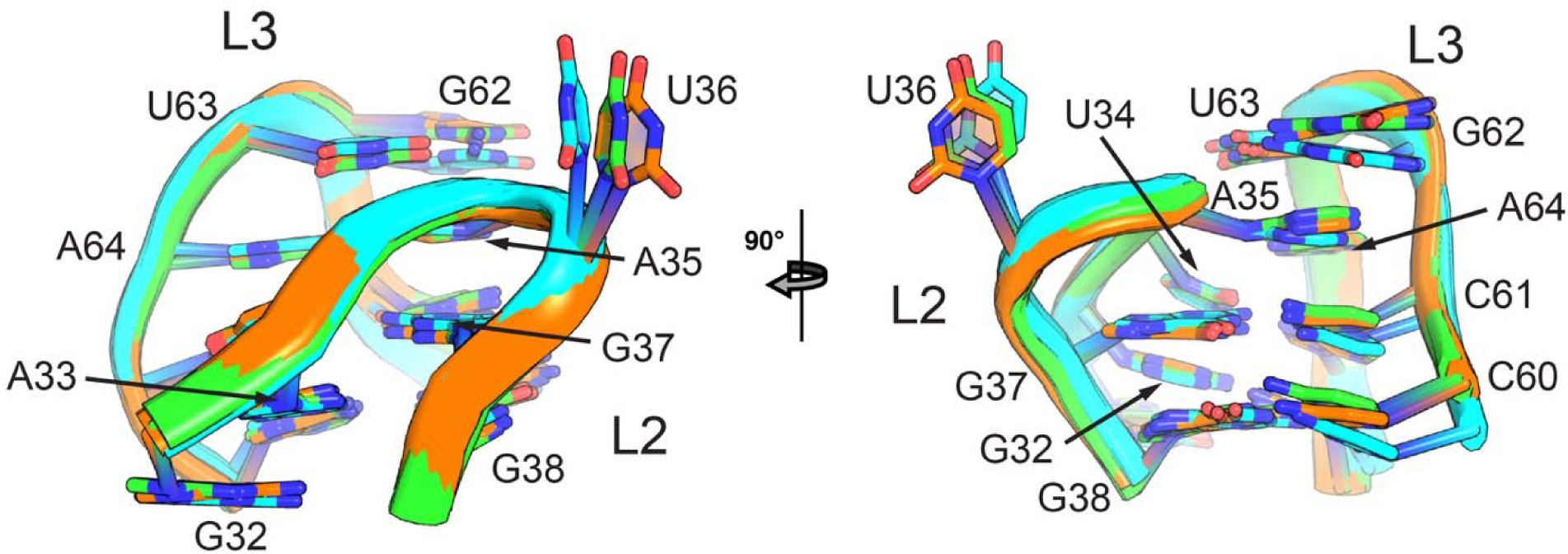
Superimposition of the L2-L3 interaction. The L2-L3 region of the parental *B. subtilis xpt* guanine riboswitch aptamer domain (green), the DGR-1B aptamer (orange) and DGR-1A aptamer (cyan) are superimposed upon one another to show that the long-range tertiary interaction of the parental RNA is preserved in the two selected aptamers.

### The scaffold is reorganized by a large deletion acquired during selection

Although the global fold is preserved, substantial secondary structure rearrangements occur in regions not explicitly randomized during the selection. The most pronounced change is a deletion of five or six nucleotides from the 3’-side of the constant region of the P1 helix in both DGR-1A and DGR-1B (red box, **Supplementary Figure S6**). Consequently, nucleotides originally forming the P1 helix are incorporated into the adjacent joining regions J1/2 and J3/1. This deletion results in the expansion of J1/2 from three nucleotides in the parental GR to thirteen nucleotides in both DGR aptamers (green strand, **Figure 2**), forming a large “S”-shaped structure that participates extensively in formation of the core of the 3WJ. During systematic evolution of ligands by exponential enrichment (SELEX), deletions within the constant region weakened the secondary structure of P1 helix, facilitating reverse transcription of the RNA and conferring a selective advantage during amplification (24).

As a result of the P1 deletion and sequence differences within J3/1, the secondary structure of P3 diverges between the two aptamers. DGR-1B preserves the P3/L3 stem-loop, including the U47-A65 Watson-Crick-Franklin (WCF) base pair derived from the randomized region. Proximal the 3WJ, the helix contains either a stacked but unpaired U46 followed by an A45-U66 WCF pair (protomer A, **Figure 2A**) or a wobble U46•U66 pair with A46 stacked but unpaired (protomer B).

In DGR-1A, additional local rearrangements are observed. The terminal A54-U62 base pair adjacent to L3 is disrupted but remains stacked, and U64 extrudes from the helix to permit formation of three sequential G-C WCF pairs. Deletion of U64 modestly increased dopamine affinity (ΔU64; K_rel_ = 0.59; **Table 1**), indicating that this bulged nucleotide slightly destabilizes the folded RNA. Proximal to the 3WJ, P3 forms a U49-A68 WCF pair from the randomized sequence as observed in DGR-1B, but this pair is preceded by a bulged uridine (U48) followed by a G47-C69 WCF pair. Despite these differences, the helical distance between the base of L3 and the 3WJ is conserved across both aptamers.

These findings demonstrate that nonrandomized scaffold regions can undergo substantial secondary structure reorganization while preserving aptamer architecture. Unlike scaffolded selection strategies that explicitly introduce insertions or deletions to further add variation to a randomized sequence library (36), the DGR aptamers illustrate that indels arising during reverse transcription can provide additional, unanticipated sources of sequence diversity that selection can act upon.

### The three-way junction is organized to form a dopamine-selective binding site

The 3WJ of the DGR aptamers is formed through an extensive network of base-base and base-backbone interactions involved all three joining strands (J1/2, J2/3, and J3/1). The base of the junction is established by a long-range interaction between C42 in J2/3 and the junction-proximal A8-U74 WCF pair of P1 (interaction *i*, **Figure 4**). Notably, this triple base pair is structurally identical to the interaction between C50 of J2/3 and the junction-proximal A21-U75 base pair in P1 that anchors the turn of J2/3 to P1 in the GR scaffold.

**Figure 4.**
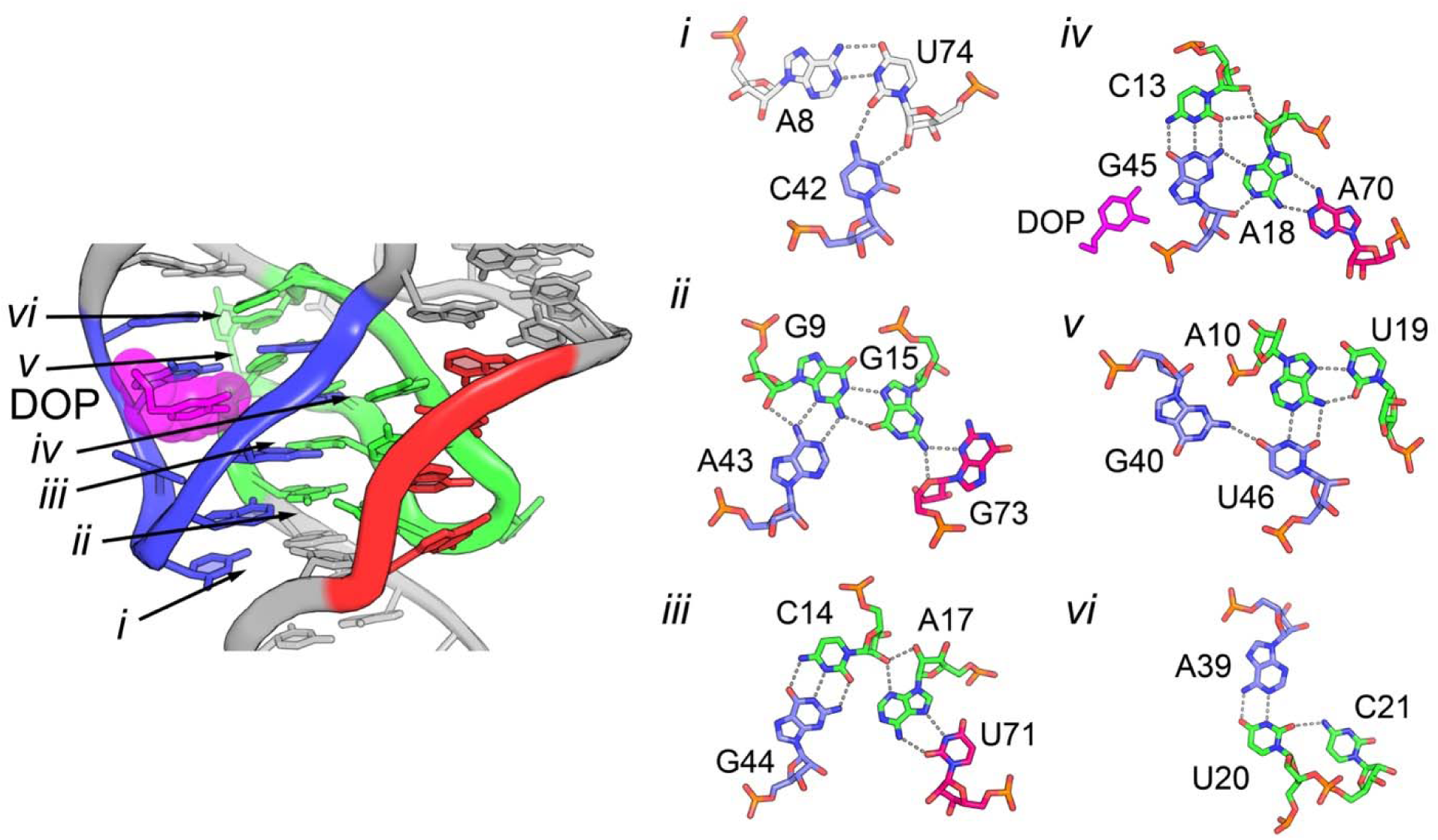
Detailed view of the three-way junction of DGR-1A. (A) Close-up cartoon representation of the junction, with the three joining strands colored as in Figure 3 and dopamine represented in magenta. Numbering *i*-*vi* denotes the positions of base-base interactions shown in panel (B). (B) Details of base-mediated interactions in the three-way junction, with dashes denoting hydrogen bonding interactions with distances >3.5 Å.

The core of the 3WJ is defined by three base quadruple interaction networks. The first (interaction *ii*, **Figure 4**) involves G9, G15, A43, and G73 and is centered on a base triple formed through interactions between the WCF edge of A43 and the sugar edge of G9 with another between the WCF of G9 and the Hoogsteen face of G15. This triple base interaction is reinforced by bifurcated hydrogen bonding interactions between U41(O2’) and A43(N7)/G9(O2’) and between G15(N2) and G73(N3)/G73(O4’). Mutation of G73 to A, which would preserve its interactions with G15, does not impact dopamine binding (**Table 1**). The second quadruple consists of two base pairs connected by a ribose zipper interaction (interaction *iii*, **Figure 4**): a C14-G44 WCF pair and a *trans* Hoogsteen A17-U71 pair. The ribose zipper interaction is formed by bifurcated hydrogen bonds between the 2’-hydroxyl group of C14 and A17(N3/O2’).

The third interaction network (interaction *iv*, **Figure 4**) involves C13, A18, G45, and A70 and consists of a C13-G45 WCF pair and a *trans* Hoogsteen A18-WCF A70 pair. These pairs are linked via a type-I A-minor triple interaction involving the sugar edge of A18 and the C13-G45 base pair. Mutation of G45 to A eliminates detectable dopamine binding while the compensatory mutation C13U/G45A only weakly restores ligand binding to ∼5.4 mM (**Table 1**). These results indicate that the C13-G45 WCF pair is critically important. Since the C13U/G45A mutation maintains the potential to fully interact with dopamine since the potential hydrogen bonding pattern to dopamine is preserved (see next section for details of ligand-RNA interactions), it is likely that the identity of C13-G45 is necessary for proper folding of the 3WJ. Three junction nucleotides (C11, U12, and U16) are bulged and unstacked; mutation of U12 to A does not alter dopamine binding, consistent with its peripheral role in the structure (**Table 1**).

In addition to providing a key ligand-interacting nucleotide, the C13-G45/A18•A70 platform supports coaxial stacking of both the P2 and P3 helices onto the 3WJ, supporting their side-by-side orientation. The interface between the 3WJ and P2 is composed of a base quadruple and a base triple. The quadruple is comprised of a *trans* A10-U46 WCF interaction and a *trans* Hoogsteen A10•U19 pair (interaction *v*, **Figure 4**). A single hydrogen bond exists between G40(N2) and U46(O4) with G40 also serving as the ceiling of the dopamine binding pocket. The base triple involves a U20-A39 WCF pair interacting with C21 through a single hydrogen bond (interaction *vi*, **Figure 4**). The side-by-side arrangement of the sequential U20 and C21 nucleotides is similar to that of the A-platform motif. Stacked upon these interactions is a G22-C38 WCF pair, conserved from the GR scaffold. The sequence and local structure of the 3WJ-P2 interface is identical in apo DGR-1B and ligand-bound DGR-1A.

In contrast, the 3WJ-P3 interface differs between the two aptamers. In DGR-1A, five nucleotides whose positions were randomized in the original selection library form this interface, including a G47-C69 WCF pair followed by a bulged U48 and a U49-A68 WCF pair. Deletion of U48 had no effect on dopamine binding affinity (**Table 1**), and deletion of both bulged P3 nucleotides (ΔU48/ΔU64) slightly increased ligand affinity (**Table 1**) indicating that these residues do not contribute productively to aptamer function. In DGR-1B, the equivalent region forms a U45-U66 WCF pair followed by a stacked but unpaired U46 and a U47-A65 pair.

### Ligand binding is mediated by a cation

A distinct feature of both apo and dopamine-bound aptamers is the presence of a cation in a deep crevice in the binding pocket within J2/3. In the apo structure of DGR-1B, a strong electron density peak (∼4.5σ) is observed at this site (**Figure 5A**). This density lies within 3 Å of G42(O6), G43(O6), and a nonbridging phosphate oxygen of A8 consistent with a bound cation. Based upon the peak height, coordination geometry, and the cations in the crystallization condition (Sr^2+^ and Li^+^), this density was assigned as a strontium ion.

**Figure 5.**
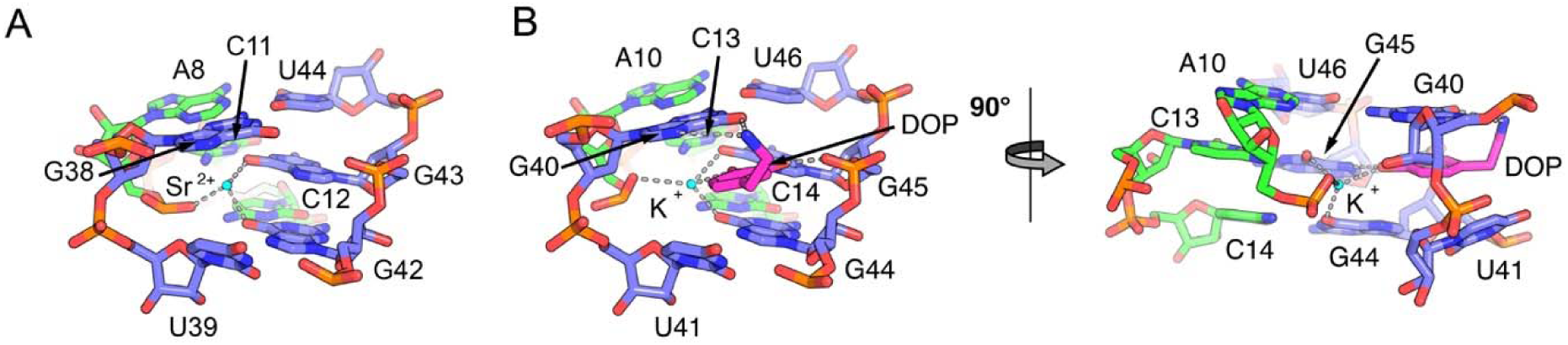
Metal-mediated interactions between dopamine and the DGR-1A aptamer. (A) Detailed view of the region of J2/3 (blue) and J1/2 (green) in the unbound DGR-1B aptamer that harbors a deep pocket with a strontium ion (cyan) situated adjacent to A8, G42, and G43. Dashed grey lines represent RNA-metal distances ≤3.5 Å. (B) The same view and a 90° rotation of the DGR-1A aptamer with dopamine bound. A potassium ion (cyan) is positioned in nearly the identical spot as the Sr^2+^ ion in DGR-1B. The 4-position hydroxyl group of dopamine makes an additional contact to the metal cation.

In the dopamine-bound DGR-1A structure, a similarly positioned electron density peak is observed adjacent to the equivalent oxygens: G44(O6), G45(O6), and A10(OP1) (**Figure 5B**). In addition, the O2 of the dopamine catechol ring is near the electron density peak. Given the peak characteristics and the crystallization conditions (K^+^ and Mg^2+^), this peak was assigned as a potassium ion.

To assess the functional importance of monovalent cation identity, we measured dopamine binding to DGR-1A in the presence of different monovalent cations (**Table 2**). The highest affinity was observed in the presence of K^+^, which was the dominant monovalent cation used during the selection (24). Sodium supported dopamine binding but with approximately threefold lower affinity whereas Li^+^ and Cs^+^ reduced affinity by an approximately 20-fold decrease. SHAPE probing confirmed that monovalent cation identity did not significantly affect global or local RNA folding (**Supplementary Figure S7**), indicating that the observed affinity differences arise from direct involvement of the cation in ligand recognition.

**Table 2:**
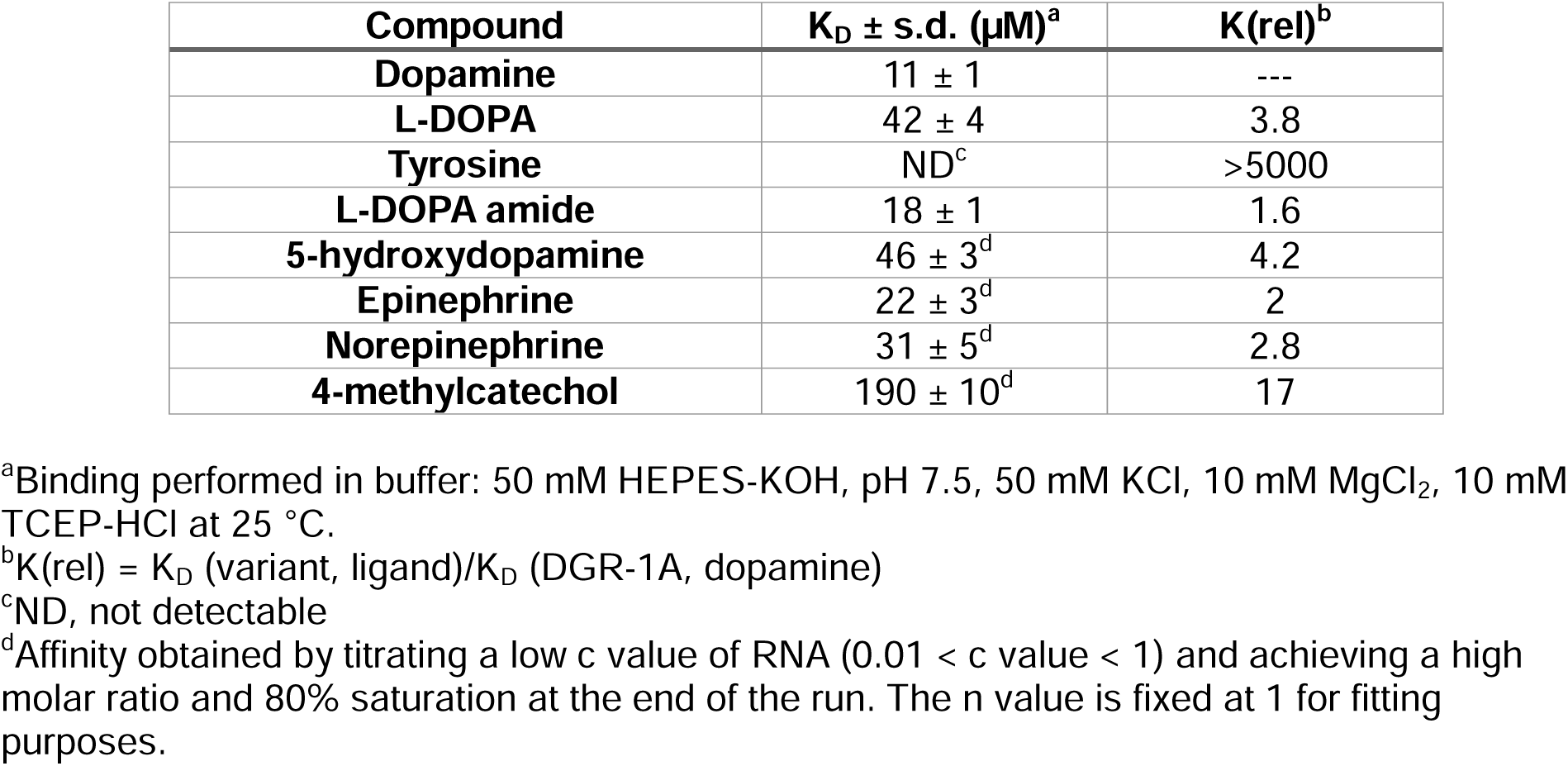
DGR-1A binding to dopamine analogs.

| Compound | $K_D \pm \text{s.d. } (\mu\text{M})^a$ | $K(\text{rel})^b$ |
| --- | --- | --- |
| **Dopamine** | $11 \pm 1$ | --- |
| **L-DOPA** | $42 \pm 4$ | 3.8 |
| **Tyrosine** | ND<sup>c</sup> | >5000 |
| **L-DOPA amide** | $18 \pm 1$ | 1.6 |
| **5-hydroxydopamine** | $46 \pm 3^d$ | 4.2 |
| **Epinephrine** | $22 \pm 3^d$ | 2 |
| **Norepinephrine** | $31 \pm 5^d$ | 2.8 |
| **4-methylcatechol** | $190 \pm 10^d$ | 17 |

<sup>a</sup>Binding performed in buffer: 50 mM HEPES-KOH, pH 7.5, 50 mM KCl, 10 mM MgCl<sub>2</sub>, 10 mM TCEP-HCl at 25 °C.
<sup>b</sup> $K(\text{rel}) = K_D (\text{variant, ligand})/K_D (\text{DGR-1A, dopamine})$
<sup>c</sup>ND, not detectable
<sup>d</sup>Affinity obtained by titrating a low c value of RNA ( $0.01 < c \text{ value} < 1$ ) and achieving a high molar ratio and 80% saturation at the end of the run. The n value is fixed at 1 for fitting purposes.

Cation-mediated ligand recognition is a recurring feature of riboswitches. For example, the *Thermotoga maritima* lysine riboswitch uses a single K^+^ cation mediating recognition of the lysine carboxylate group and displays a strong preference for K^+^ over Na^+^ (72). Similar ion-mediated recognition has been reported for the ZTP, TPP, and FMN riboswitches (16,73–79), underscoring this mechanism as a common RNA strategy for binding oxygen-bearing moieties in ligands.

### The DGR aptamer primarily recognizes the catechol moiety of dopamine

The ligand binding pocket is defined primarily by the J2/3 strand that forms a loop structure. The loop adopts a canonical U-turn motif with the UNR consensus sequence (80) comprised of U41, C42, and A43 forming the turn. Within this motif, the first uridine residue anchors the turn via a hydrogen bond between N3 and G44(OP2) while the second and third nucleotide bases stack upon each other on the other side of the tight turn.

On the 3’-side of the loop, two purine residues (G44 and G45) form an open pocket for ligand binding and are anchored to the 3WJ core via WCF base pairs with C13 and C14 of J1/2. When G44 is mutated to a C, detectable dopamine binding is eliminated. The compensatory mutation, C14G/G44C, restores binding to micromolar affinity (**Table 1**), highlighting their essential structural role. The J2/3 loop is closed by a base triple interaction between A10, G40, and U46 where A10 and U46 form a WCF pair and G40 and U46 form a single hydrogen bonding interaction between G40(N2) and U46(O2). Although superficially this arrangement is reminiscent of a T-loop motif, it is structurally distinct (81).

The dopamine binding pocket is centered within the J2/3 turn. The ceiling of the pocket is formed by G40/U46 with G40(O6) within hydrogen bonding distance of the dopamine amine (**Figure 5B**). The catechol ring lies adjacent to G45 with dopamine O1 forming a hydrogen bond with G45(OP2) and dopamine (O2) interacting with G45(N7). The floor of the binding pocket is formed by nucleotides U41 and G44. Aromatic stacking interactions between dopamine and G40 and G44 is present but modest while significant open space adjacent to the C5/C6 positions of the catechol ring suggests that the pocket is not optimized for maximal shape complementarity.

To assess the contributions of functional groups, we measured the binding of dopamine analogs to DGR-1A. The original selection of the DGR-1A and DGR-1B aptamers used L-DOPA immobilized to a column through the carboxylate group such that this functional group was converted into an amide that connected to a flexible linker (24). Both L-DOPA and L-DOPA amide bind more weakly than dopamine, indicating that neither the carboxylate group nor its conversion to an amide contributes productively to binding (**Table 3**). DGR-1A also showed limited discrimination between dopamine and its derivatives epinephrine and norepinephrine. Indicative of the dopamine amine contributing to binding, conversion of the dopamine ethylamine group to a methyl group in 4-methylcatechol reduced affinity by 17-fold. In contrast, removal of a single hydroxyl group in L-DOPA to form tyrosine abolished binding, supporting recognition of the catechol ring. Addition of a hydroxyl group at the 5-position resulted in an affinity ∼4-fold weaker than dopamine, consistent with the open space observed in this region being able to accommodate the additional moiety. Together, these data demonstrate that the DGR-1 aptamers primarily recognize the catechol group of dopamine with additional recognition of the dopamine amine.

**Table 3:** Monovalent cation dependence of dopamine binding to DGR-1A.

| **Cation** | **$K_D \pm \text{s.d. } (\mu\text{M})^a$** |
| --- | --- |
| **Li<sup>+</sup>** | $110 \pm 10^b$ |
| **Na<sup>+</sup>** | $17 \pm 4^b$ |
| **K<sup>+</sup>** | $6 \pm 3^b$ |
| **Cs<sup>+</sup>** | $130 \pm 10^b$ |

<sup>a</sup>Binding performed in buffer: 50 mM HEPES acid, pH 7.5, 125 mM monovalent cation, 10 mM MgCl<sub>2</sub>, 10 mM TCEP-HCl at 25 °C. Buffer was adjusted to the correct pH with the appropriate cation hydroxide and the chloride salt was added to obtain a total of 125 mM of the monovalent cation.
<sup>b</sup>Affinity obtained by titrating a low c value of RNA ( $0.01 < c \text{ value} < 1$ ) and achieving a high molar ratio and 80% saturation at the end of the run. The n value is fixed at 1 for fitting purposes.

### The aptamer undergoes minimal conformational change upon ligand binding

Comparison of the apo DGR-1A and dopamine-bound DGR-1B structures suggests that the RNA is largely preorganized for ligand binding. Because apo riboswitch structures often crystallize in conformations similar to the bound state that does not reflect its behavior in solution, we probed local RNA flexibility using SHAPE chemical probing which reports on local backbone dynamics via 2’-hydroxyl reactivity (82).

SHAPE analysis of DGR-1A in the presence and absence of dopamine revealed that the aptamer is predominantly preorganized. In the absence of dopamine, reactivity is observed primarily at bulged nucleotides (nucleotides 11, 12, 16, 48, 64, 72, and 73) which are expected to be flexible (**Figure 6A & B, Supplementary Figure S8**). Strong reactivity at nucleotides 57 and 58 corresponds to the L3 turn, matching reactivity patterns observed in the parental GR scaffold and supporting conservation of the L2-L3 interaction (60). Moderate reactivity in nucleotides 39-41 near the binding site decreased weakly upon dopamine binding, indicating limited local rearrangement.

**Figure 6.**
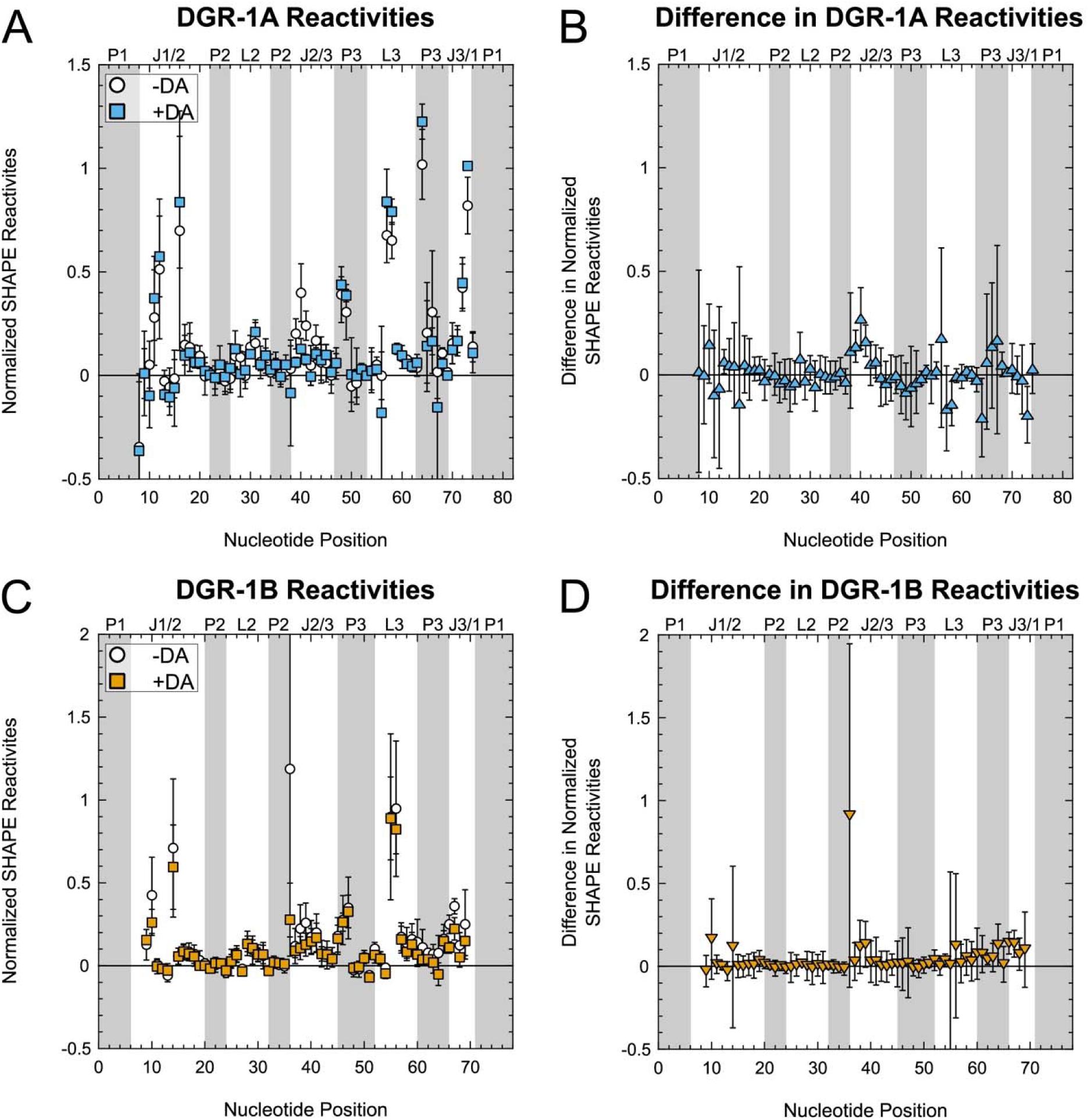
SHAPE chemical probing of DGR aptamers in the absence and presence of dopamine. RNAs were probed with 1M7 with or without 1 mM dopamine (A, C). Positive values indicate greater reactivity. The differences in SHAPE reactivities were determined by subtracting the intensities of the apo state from the bound state (B, D). Greater values indicate differences in reactivity upon ligand binding. Both DGR-1A and DGR-1B show little to no change in SHAPE reactivities between the apo and bound states indicated by differences close to 0.

DGR-1B exhibited similar SHAPE profiles with fewer reactive positions overall (**Figure 6C & D, Supplementary Figure S9**). Bulged nucleotides 10 and 14 and nucleotides 55 and 56 at the L3 turn showed strong reactivity that was unchanged upon ligand binding. Nucleotides in J2/3 show low reactivity with very slight protection in nucleotides 38 and 39 upon dopamine binding. Together, these data support a predominantly preorganized, lock-and-key-like mechanism of ligand recognition by the DGR-1 aptamers.

## CONCLUSIONS

A critical challenge in the development of RNA aptamers is that despite decades of *in vitro* selection experiments raising hundreds of small molecule binding aptamers, most aptamers have failed to function robustly across diverse biological and technological contexts. In contrast, naturally occurring riboswitches rely on evolutionarily optimized aptamer domains that fold rapidly and reproducibly under stringent co-transcriptional constraints. Scaffolded selection was developed to bridge this gap by embedding new ligand-binding pockets within the highly stable architectures of biological riboswitches (24). The structural analysis presented in this study provides further evidence that this strategy succeeds in enforcing global fold conservation while accommodating locally diverse pockets in the 3WJ required for new ligand specificity.

Across all structurally characterized aptamers generated by scaffolded selection like the 5HTP aptamer (24), Tonic (38), Squash (37), and the DGR aptamers, the defining tertiary hallmarks of the parental scaffold are preserved. The DGR aptamers extend this paradigm by demonstrating that regardless of local deviations such as disruption of P1 coaxial stacking with P3, the fundamental scaffold architecture remains intact and capable of hosting a selective ligand binding pocket. These findings reinforce the central hypothesis motivating scaffolded selection: leveraging riboswitch-derived “superfolder” architectures can systematically bias selection toward aptamers with robust folding and functional resilience.

Importantly, the DGR aptamers also reveal that scaffolded selection does not merely preserve static features of the parent fold but enables productive structural plasticity. Deletions acquired during SELEX and subsequent remodeling of the 3WJ generated a selective dopamine binding pocket that relies on cation-mediated ligand recognition while maintaining a largely pre-organized RNA conformation. This balance between architectural conservation and local adaptability highlights how a single RNA fold can host divergent binding pockets and recognize chemically distinct ligands, a property long observed in natural riboswitch families but rarely achieved with synthetic aptamers.

Finally, the successful deployment of the DGR-1B aptamer as the sensing domain of a functional dopamine-responsive riboswitch underscores the practical relevance of this approach (83). In this design, the DGR-1A and DGR-1B aptamers, amongst a set of other dopamine aptamers, were attached to an expression platform containing an intrinsic terminator that invades into P1 and directs RNA polymerase to abort transcription in the absence of L-DOPA/dopamine. While both the DGR-1A and DGR-1B (RS1 and RS2 riboswitches, respectively, in (83)) both demonstrated dopamine-responsiveness, DGR-1B outperformed DGR-1A and performed better than a riboswitch designed using a dopamine aptamer from a prior deep selection of dopamine aptamers (80 sequential randomized nucleotides in starting pool). The RS2 riboswitch in a cell-free system detected low concentrations of dopamine (1 – 3 µM) spiked into human urine, which outperformed an established ELISA (83). While other issues with the riboswitch assay prevented its further application, these results establish scaffolded selection as a powerful framework for generating small-molecule RNA aptamers that combine the adaptability of *in vitro* selection with the folding fidelity and functional robustness characteristic of natural riboswitches.

## Data Availability

Atomic coordinates and structure factor amplitudes have been deposited into the Protein Data Bank (PDB) database under accession codes 12CI (DGR-1A, bound), 12CJ (DGR-1B, apo).

## Supplementary data

Supplementary data is available at NAR online.

## Acknowledgements

We thank Dr. Jay Nix and the staff of beamline 8.2.1 of the Advanced Light Source, Lawrence Berkeley National Laboratory, for their support with remote crystallographic data collection. This research used resources at Beamline 8.2.1 of the Advanced Light Source, which is a U.S. DOE Office of Science User Facility under Contract No. DE-AC02-05CH11231. Crystallography time is supported in part by the ALS-ENABLE program, funded by the National Institutes of Health, National Institute of General Medical Sciences (grant P30 GM124169). We thank the Macromolecular X-ray Crystallography Core (RRID:SCR_019310) at the University of Colorado Boulder for crystallographic data collection and the Shared Instruments Pool (RRID: SCR_018986). The Typhoon 5 is funded by NIH Shared Instrumentation Grant S10OD034218-01. We thank Dr. Annette Erbse for her assistance with the resources utilized at the University of Colorado Boulder.

## Author contributions

Conceptualization (R.T.B.), Investigation (S.H.S., D.S., S.D.H.), Formal analysis (S.H.S., D.S., S.D.H., R.T.B.), Visualization (S.H.S., R.T.B.), Resources (R.T.B.), Writing — Original Draft Preparation (R.T.B.), Writing — Review and Editing (S.H.S., R.T.B.), Funding Acquisition (R.T.B.), Supervision (R.T.B.).

