## Supplementary Information for "Structure of a dopamine-binding RNA aptamer reveals metal-mediated ligand recognition"

Supplementary Table S1. Oligonucleotide sequences used in this work.

| **Name** | **Used for** | **Sequence 5’🡪3’** |
| --- | --- | --- |
| 5’ Primer | All preps of RNA | GCGCGCGAATTCTAATACGACTCACTATAG |
| DGR-1A P1-8 | Crystallography, ITC, SHAPE | CCGTAATACGACTCACTATAGGACGATAGACTCCGTAATTCGCGTGGATATGGCACGCAGTCAGGTGTTGGGCACCGTAAATGTCCCACATAGTATCGTCCA |
| 3’ Primer DGR-1A P1-8 | Crystallography | mUmGGACGATACTATGTGGGACATTTACGG |
| 3’ Primer DGR-1A P1-8 | ITC | TGGACGATACTATGTGGGACATTTACGG |
| 3’ Primer DGR-1A P1-8 + SHAPE cassette | SHAPE | GAACCGGACCGAAGCCCGTGGACGATACTATGTGGGACATTTACGG |
| DGR-1B P1-6 | Crystallography, ITC, SHAPE | CCGTAATACGACTCACTATAGGACTAGACTCCGTAATTCGCGTGGATATGGCACGCAGTCAGGTGTTGGGCACCGTAAATGTCCCACATAGTAGTCCA |
| 3’ Primer DGR-1B P1-6 | Crystallography, ITC | mUmGGACTACTATGTGGGACATTTACGG |
| 3’ Primer DGR-1B P1-6 + SHAPE cassette | SHAPE | GAACCGGACCGAAGCCCGTGGACTACTATGTG GGACATTTACGG |

Supplementary Table S2. Mutant DGR-1A oligonucleotides used for recursive PCR. Changes to the wildtype sequence are highlighted in yellow. Nucleotides flanking deletions to the wildtype sequence are highlighted in cyan.

| **Mutant** | **Sequence 5’ 🡪3’** |
| --- | --- |
| 5’ Primer used for all mutants | GCGCGCGAATTCTAATACGACTCACTATAG |
| **G45A** | |
| 5’ Fragment WT | CCGTAATACGACTCACTATAGGACGATAGACTCCGTAATTCGCGTGGATATGGCACGCAG |
| 3’ Fragment G45A | GGACGATACTATGTGGGACATTTACGGTGCCCAACATCTGACTGCGTGCCATATCCACGC |
| 3’ Primer WT | TGGACGATACTATGTGGGACATTTACGG |
| **C13U/G45A** | |
| 5’ Fragment C13U | CCGTAATACGACTCACTATAGGACGATAGACTTCGTAATTCGCGTGGATATGGCACGCAG |
| 3’ Fragment G45A | GGACGATACTATGTGGGACATTTACGGTGCCCAACATCTGACTGCGTGCCATATCCACGC |
| 3’ Primer WT | TGGACGATACTATGTGGGACATTTACGG |
| **G44C** | |
| 5’ Fragment WT | CCGTAATACGACTCACTATAGGACGATAGACTCCGTAATTCGCGTGGATATGGCACGCAG |
| 3’ Fragment G44C | GGACGATACTATGTGGGACATTTACGGTGCCCAACACGTGACTGCGTGCCATATCCACGC |
| 3’ Primer WT | TGGACGATACTATGTGGGACATTTACGG |
| **C14G/G44C** | |
| 5’ Fragment C14G | CCGTAATACGACTCACTATAGGACGATAGACTCGGTAATTCGCGTGGATATGGCACGCAG |
| 3’ Fragment G44C | GGACGATACTATGTGGGACATTTACGGTGCCCAACACGTGACTGCGTGCCATATCCACGC |
| 3’ Primer WT | TGGACGATACTATGTGGGACATTTACGG |
| **U12A** | |
| 5’ Fragment U12A | CCGTAATACGACTCACTATAGGACGATAGACACCGTAATTCGCGTGGATATGGCACGCAG |
| 3’ Fragment WT | GGACGATACTATGTGGGACATTTACGGTGCCCAACACCTGACTGCGTGCCATATCCACGC |
| 3’ Primer WT | TGGACGATACTATGTGGGACATTTACGG |
| **G73A** | |
| 5’ Fragment WT | CCGTAATACGACTCACTATAGGACGATAGACTCCGTAATTCGCGTGGATATGGCACGCAG |
| 3’ Fragment G73A | GGACGATATTATGTGGGACATTTACGGTGCCCAACACCTGACTGCGTGCCATATCCACGC |
| 3’ Primer G73A | GGACGATATTATGTGGGACATTTACGG |
| **ΔU48** | |
| 5’ Fragment WT | CCGTAATACGACTCACTATAGGACGATAGACTCCGTAATTCGCGTGGATATGGCACGCAG |
| 3’ Fragment ΔU48 | GGACGATACTATGTGGGACATTTACGGTGCCCACACCTGACTGCGTGCCATATCCACGC |
| 3’ Primer WT | TGGACGATACTATGTGGGACATTTACGG |
| **ΔU64** | |
| 5’ Fragment WT | CCGTAATACGACTCACTATAGGACGATAGACTCCGTAATTCGCGTGGATATGGCACGCAG |
| 3’ Fragment ΔU64 | GGACGATACTATGTGGGCATTTACGGTGCCCAACACCTGACTGCGTGCCATATCCACGC |
| 3’ Primer ΔU64 | GGACGATACTATGTGGGCATTTACGG |
| **ΔU48/ΔU64** | |
| 5’ Fragment WT | CCGTAATACGACTCACTATAGGACGATAGACTCCGTAATTCGCGTGGATATGGCACGCAG |
| 3’ Fragment ΔU48/ΔU64 | GGACGATACTATGTGGGCATTTACGGTGCCCACACCTGACTGCGTGCCATATCCACGC |
| 3’ Primer ΔU64 | GGACGATACTATGTGGGCATTTACGG |
| **WT using recursive PCR** | |
| 5’ Fragment WT | CCGTAATACGACTCACTATAGGACGATAGACTCCGTAATTCGCGTGGATATGGCACGCAG |
| 3’ Fragment WT | GGACGATACTATGTGGGACATTTACGGTGCCCAACACCTGACTGCGTGCCATATCCACGC |
| 3’ Primer WT | TGGACGATACTATGTGGGACATTTACGG |

Supplementary Table S3**.** Data collection and model statistics.

| Crystal | *Apo DGR-1B* | *DGR-1A bound to dopamine* |
| --- | --- | --- |
| PDB ID | 12CJ | 12CI |
| **Data collection** |  |  |
| Wavelength (Å) | 0.99992 | 1.54178 |
| Resolution Range (Å)^a^ | 43.940-2.930 (3.01-2.93) | 25.00-2.50 (2.54-2.50) |
| Space Group | C 1 2 1 | P 1 2_1_ 1 |
| Unit Cell |  |  |
| a, b, c (Å) | 163.60, 35.93, 97.50 | 58.03, 59.30, 67.82 |
| α, β, γ (°) | 90.00, 120.72, 90.00 | 90.00, 106.51, 90.00 |
| Total Reflections | 78424 (5892) | 28879 (1356) |
| Unique Reflections | 10888 (802) | 15514 (756) |
| R_meas_ | 0.139 (1.485) | 0.096 (0.801) |
| R_pim_ | 0.052 (0.544) | 0.052 (0.448) |
| Multiplicity | 7.2 (7.34) | 1.8 (1.7) |
| Completeness (%) | 100.0 (99.9) | 96.2 (90.3) |
| I/σ(I) | 10.5 (1.6) | 12.73 (1.30) |
| CC_1/2_ | 0.998 (0.746) | 0.945 (0.679) |
| **Refinement** |  |  |
| Resolution (Å) | 40.46 – 2.92 (3.38 – 2.92) | 24.92 – 2.59 (2.68 – 2.59) |
| No. reflections | 10887 (1002) | 13632 (1358) |
| R_work_/R_free_ | 0.2015 (0.2776)/ 0.2480 (0.3088) | 0.2245 (0.3470) / 0.2751 (0.4007) |
| No. non-hydrogen atoms | 3347 | 3701 |
| RNA | 3306 | 3498 |
| Ligand | 0 | 22 |
| Ions | 25 | 136 |
| Water | 18 | 45 |
| B-factors |  |  |
| RNA | 61.5 | 38.9 |
| Ligand | --- | 36.1 |
| Ions | 101.0 | 51.0 |
| Water | 56.9 | 43.1 |
| r.m.s. deviations |  |  |
| Bond Length (Å) | 0.002 | 0.005 |
| Bond Angle (°) | 0.578 | 0.958 |

^a^Values in parentheses are for the highest resolution shell.

Supplementary Table S4**.** Molar extinction coefficients for compounds used in this work.

| Compound | Extinction coefficient  (ε, M^-1^cm^-1^) | Wavelength (nm) | Source |
| --- | --- | --- | --- |
| Dopamine | 2600 | 280 | This study |
| L-DOPA | 2600 | 280 | This study |
| Tyrosine | 1490 | 280 | Pace, 1995 [1] |
| L-DOPA amide | 2600 | 280 | This study |
| 5-hydroxydopamine | 750 | 269.5 | This study |
| Epinephrine | 2700 | 280 | Pugsley, 2000 [2] |
| Norepinephrine | 2600 | 279 | This study |
| 4-methylcatechol | 2200 | 280 | This study |

Supplementary Table S5**.** Complete isothermal titration calorimetry thermodynamic values for DGR-1A and DGR-1B.

| **RNA Identity** | **Ligand** | **K_D_ ± s.d. (µM)** | **N value** | **ΔH ± s.d. (kcal mol^-1^)** | **ΔS ± s.d. (cal mol^-1^ K^-1^)** | **-TΔS ± s.d. (kcal mol^-1^)** |
| --- | --- | --- | --- | --- | --- | --- |
| DGR-1A | Dopamine | 11.3 ± 0.5 | 0.795 ± 0.006 | -17.8 ± 0.1 | -37.2 ± 0.5 | 11.1 ± 0.1 |
| DGR-1A | L-DOPA | 42 ± 4 | 0.51 ± 0.02 | -20.0 ± 0.9 | -47 ± 3 | 14.0 ± 0.9 |
| DGR-1A | L-DOPA amide | 18.1 ± 0.3 | 1.21 ± 0.01 | -15.5 ± 0.1 | -30.2 ± 0.4 | 9.0 ± 0.1 |
| DGR-1B | Dopamine | 28.1 ± 0.8 | 0.76 ± 0.02 | -20.17 ± 0.02 | -46.83 ± 0.09 | 13.96 ± 0.03 |
| DGR-1B | L-DOPA | 149 ± 8 | 1.03 ± 0.03 | -20.4 ± 0.7 | -51 ± 2 | 15.1 ± 0.7 |
| DGR-1B | L-DOPA amide | 47 ± 2 | 1.34 ± 0.03 | -16.2 ± 0.3 | -35 ± 1 | 10.3 ± 0.3 |

s.d. = standard deviation

Buffer = 50 mM HEPES-KOH, pH 7.5, 50 mM KCl, 10 mM MgCl_2_, 10 mM TCEP-HCl

All measurements were conducted at 25 °C.

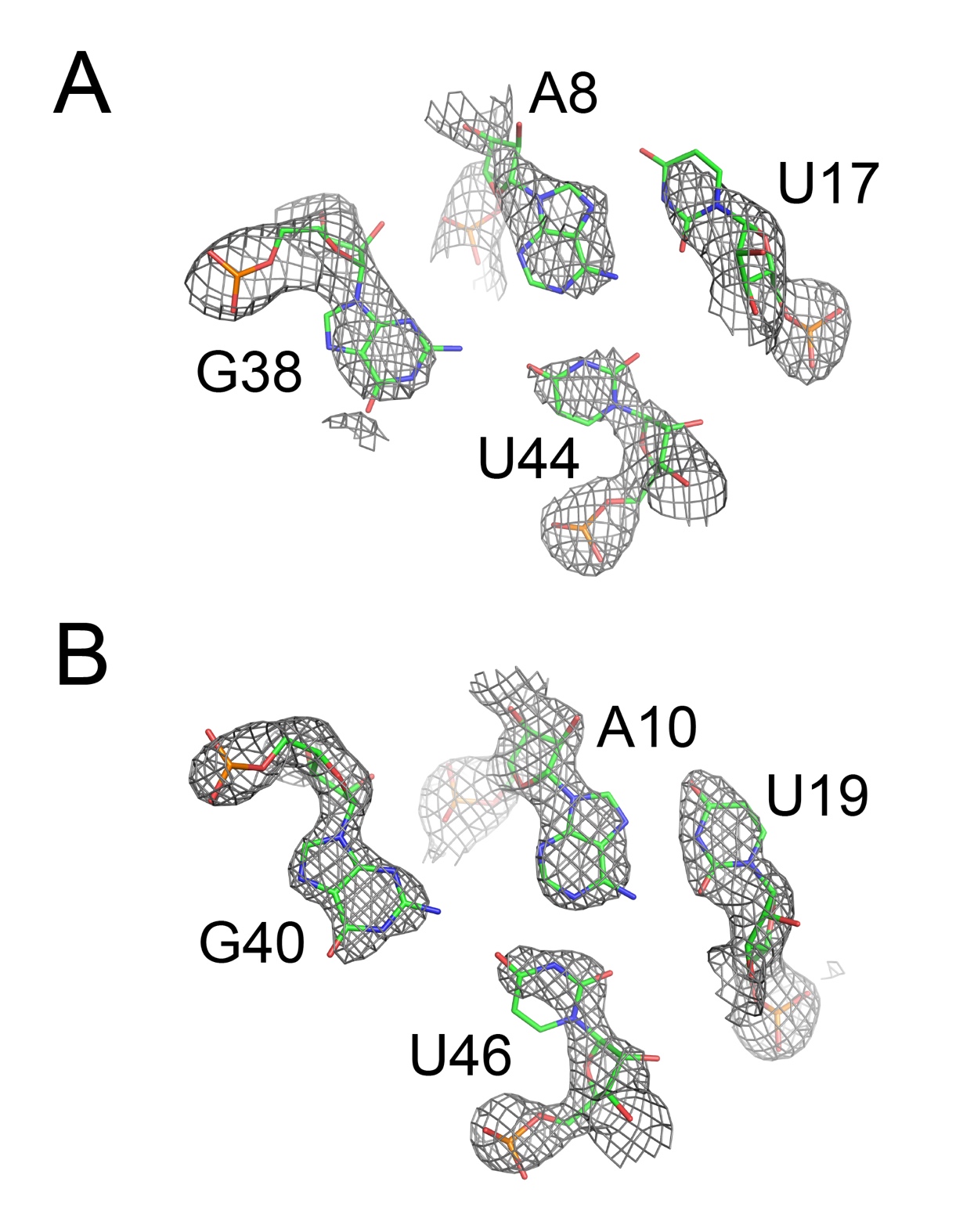

Supplementary Figure S1**.** Representative regions of composite omit 2Fo-Fc maps calculated after the final round of refinement for (A) DGR-1B (apo state) and (B) DGR-1A (bound state) aptamers. Electron density is contoured at 1.5 sigma and carved at 2.0 Å around the atoms represented by the stick model. The quartet shown is in the same orientation as pair v in Figure 4.

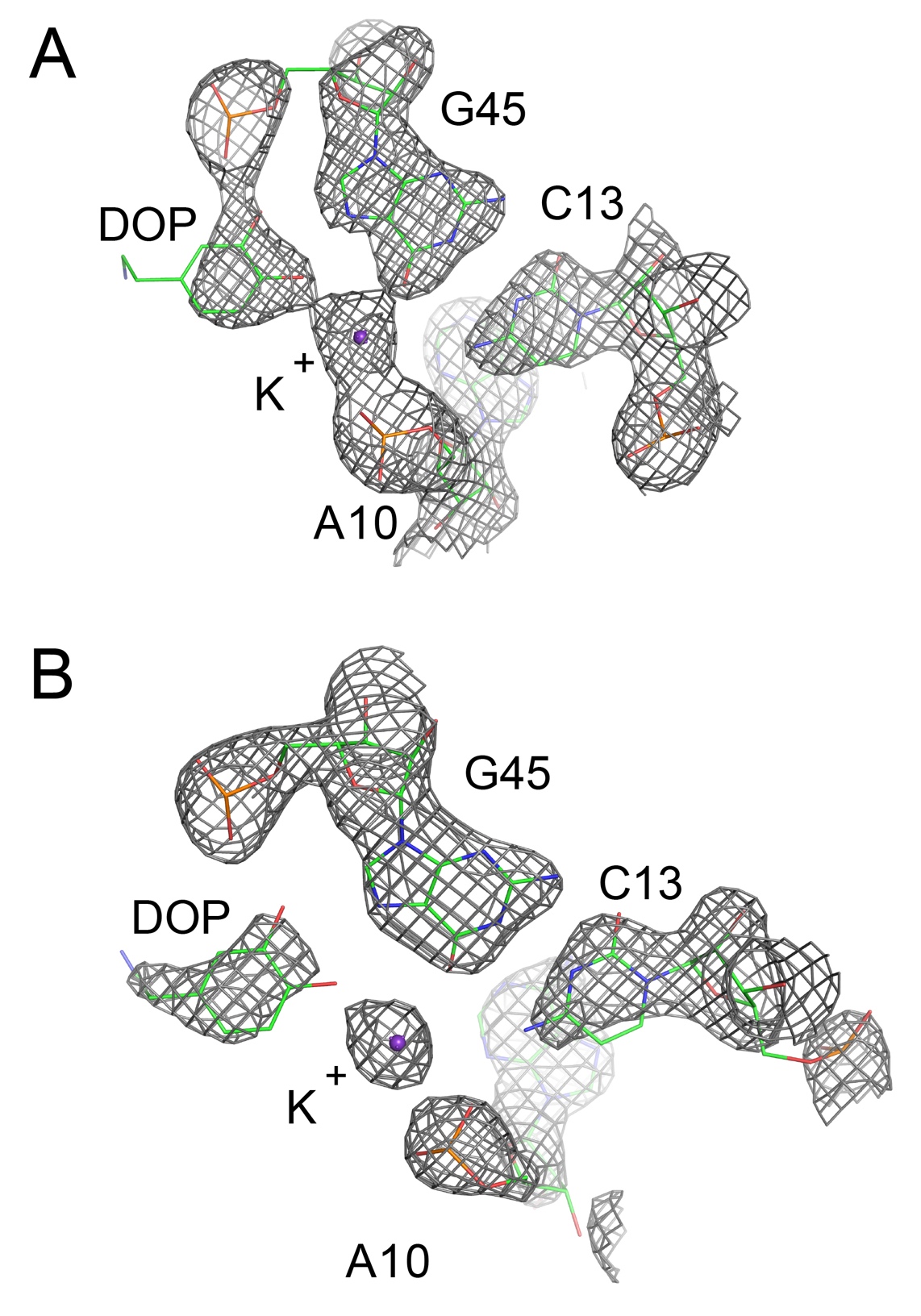

Supplementary Figure S2**.** Representative regions of composite omit 2Fo-Fc maps calculated after the final round of refinement for (A) DGR-1A (bound state) protomer A and (B) DGR-1A (bound state) protomer B. Electron density is contoured at 1.5 sigma and carved at 2.0 Å around the atoms represented by the stick model. The quartet shown is part of pair iv in Figure 4.

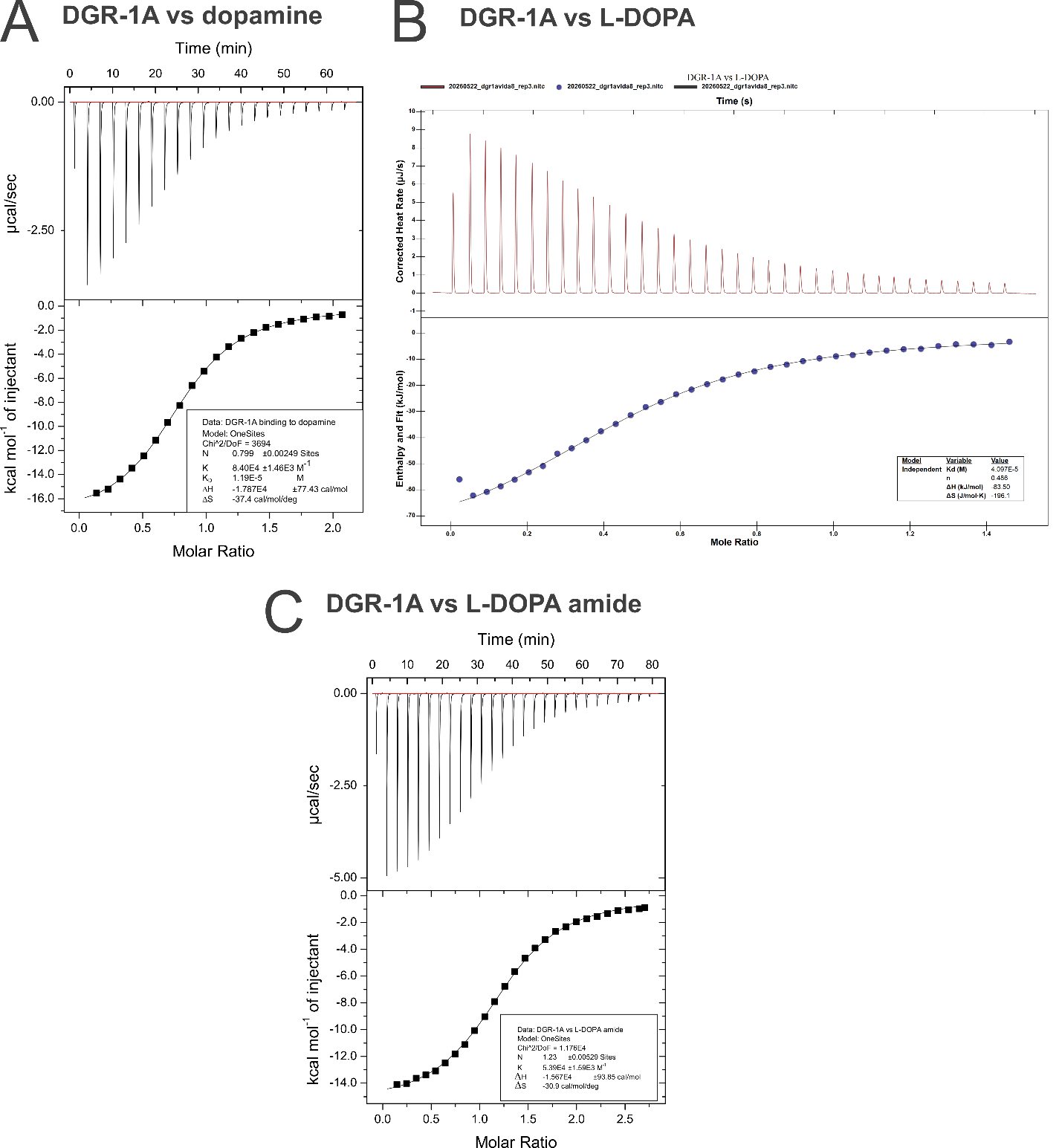

**Supplementary Figure S3.** Continued on next page.

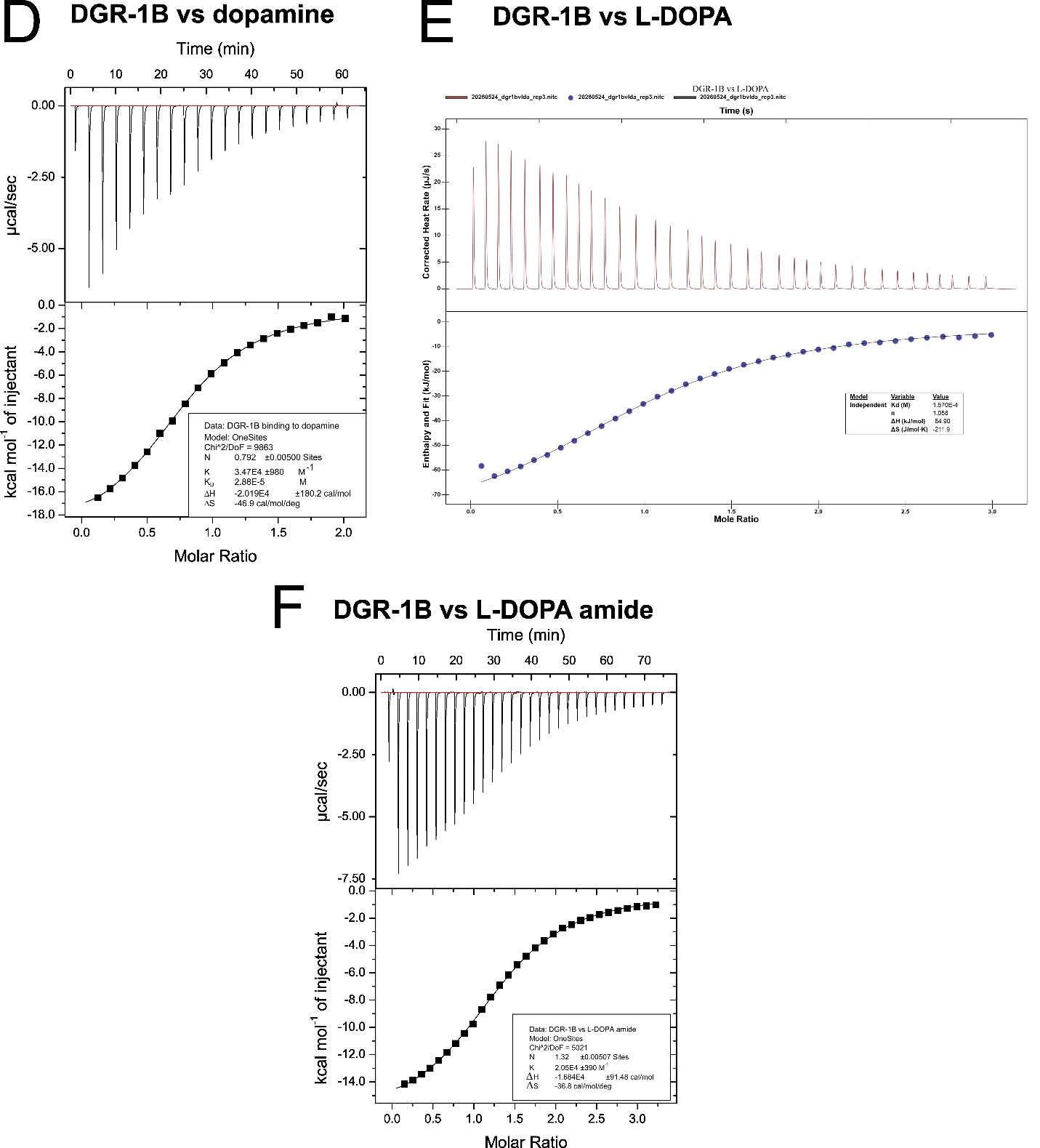

Supplementary Figure S3**.** Representative ITC thermograms at 25 °C. Exothermic binding is represented differently between the Malvern and Waters ITC instruments. Exothermic binding using the Malvern instrument results in peaks that point down while exothermic binding is represented by peaks that point upwards on the Waters instrument. (A) DGR-1A binding to dopamine. (B) DGR-1A binding to L-DOPA. (C) DGR-1A binding to L-DOPA amide. (D) DGR-1B binding to dopamine. (E) DGR-1B binding to L-DOPA. (F) DGR-1B binding to L-DOPA amide.

```
         P1-5’   J1/2     P2-5’   L2    P2-3’  J2/3   P3-5’  L3     P3-3’     P1-3’
GR       GAUAUAA........UCGCGUGGAUAUGGCACGCAAGUUUCUACCGGGCACCGUAAAUGUC.CG...ACUAUC
DGR-1A   GAUAGACUCCGUAAUUCGCGUGGAUAUGGCACGCAGUCAGGUGUUGGGCACCGUAAAUGUCCCACAUAGUAUC
DGR-1B   ACUAGACUCCGUAAUUCGCGUGGAUAUGGCACGCAGUAAGGUAUUGGGCACCGUAAAUGUC.CAUAUGUUACU
```

Supplementary Figure S4**.** Alignment of parental *B. subtilis* *xpt-pbuX* guanine riboswitch aptamer domain (GR) sequence with the DGR-1A and DGR-1B sequences. Cyan highlighted sequences denote the paired (P) regions and green denotes the terminal loops L2 and L3.

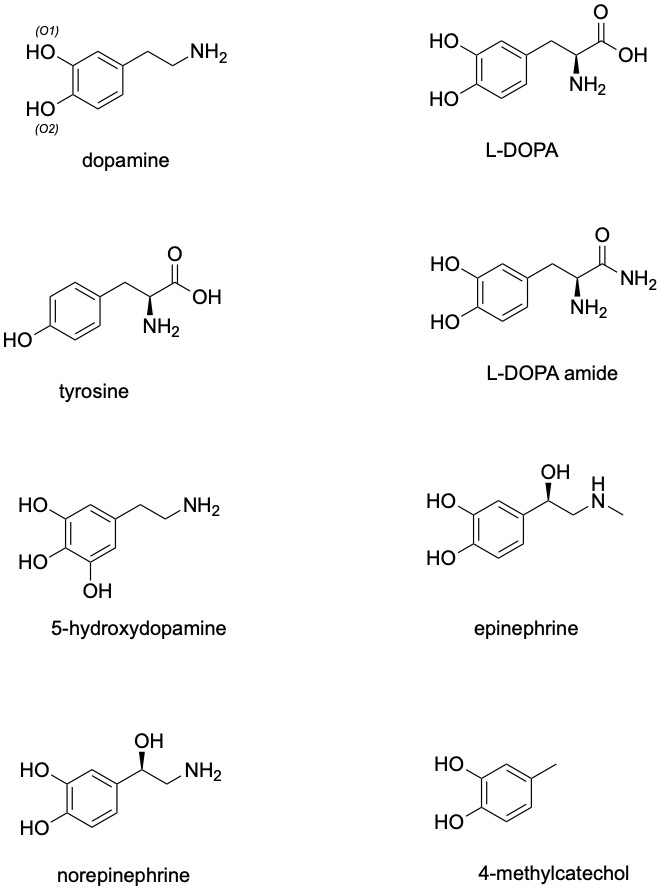

Supplementary Figure S5**.** Chemical structures of compounds used in this study.

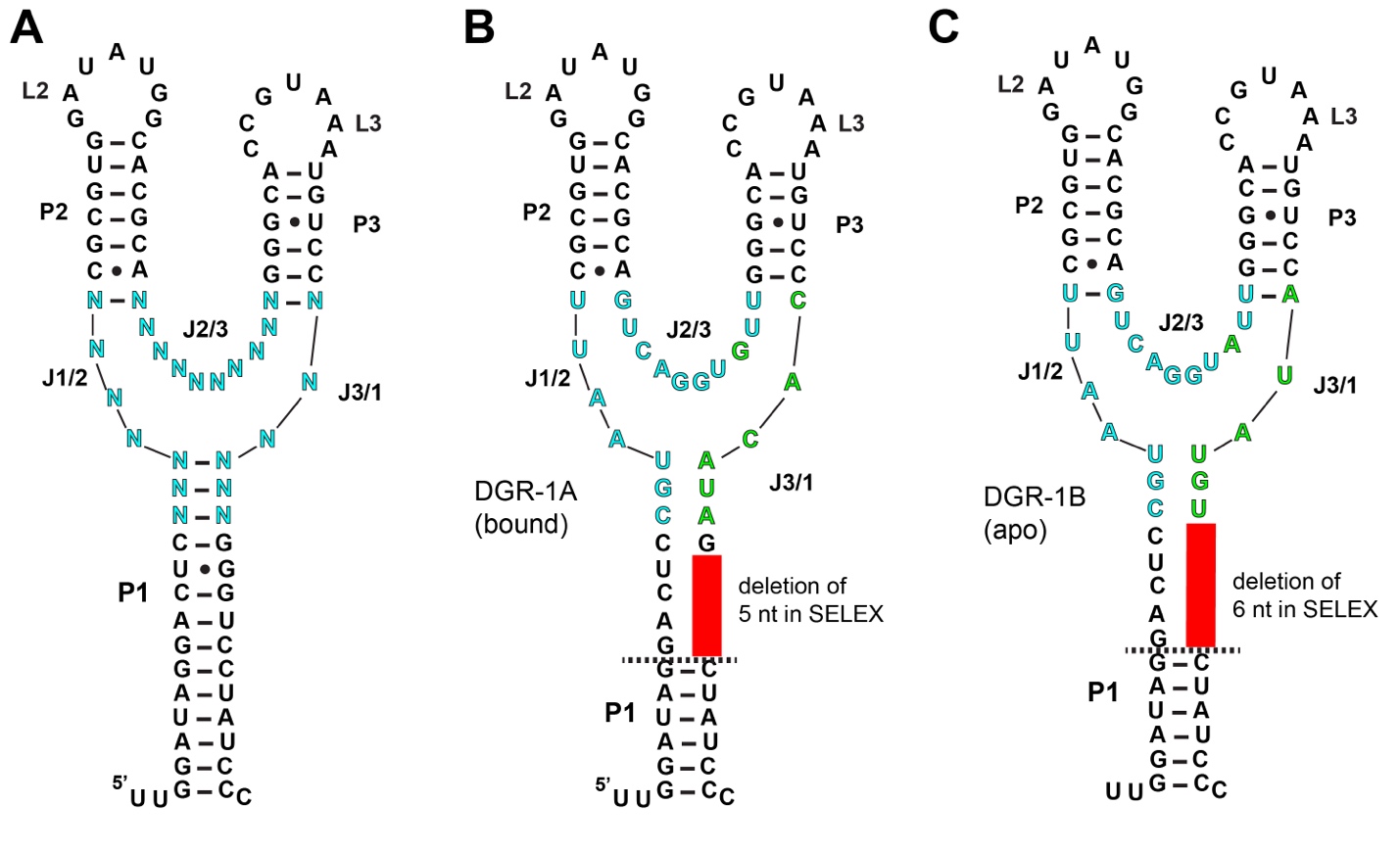

Supplementary Figure S6**.** Secondary structures of the (A) original GR scaffold library, (B) DGR-1A and (C) DGR-1B emphasizing the deletion that occurred during the selection of the dopamine aptamers as originally predicted by R2R following selection [3]. In the original selection, the library contained 23 randomized nucleotides (cyan) defining the three joining regions: J1/2, J2/3, and J3/1. During selection, a 5 or 6 nucleotide region was deleted from the 3’-side of P1 (red), enabling the residual junction-proximal nucleotides in P1 to be consumed into J1/2 and J3/1; the dashed line represents the new boundary of P1 in the DGR-1A and DGR-1B aptamers. Identities of randomized nucleotides in DGR-1A and DGR-1B are shown in cyan and green with differences between the two DGR aptamers in green.

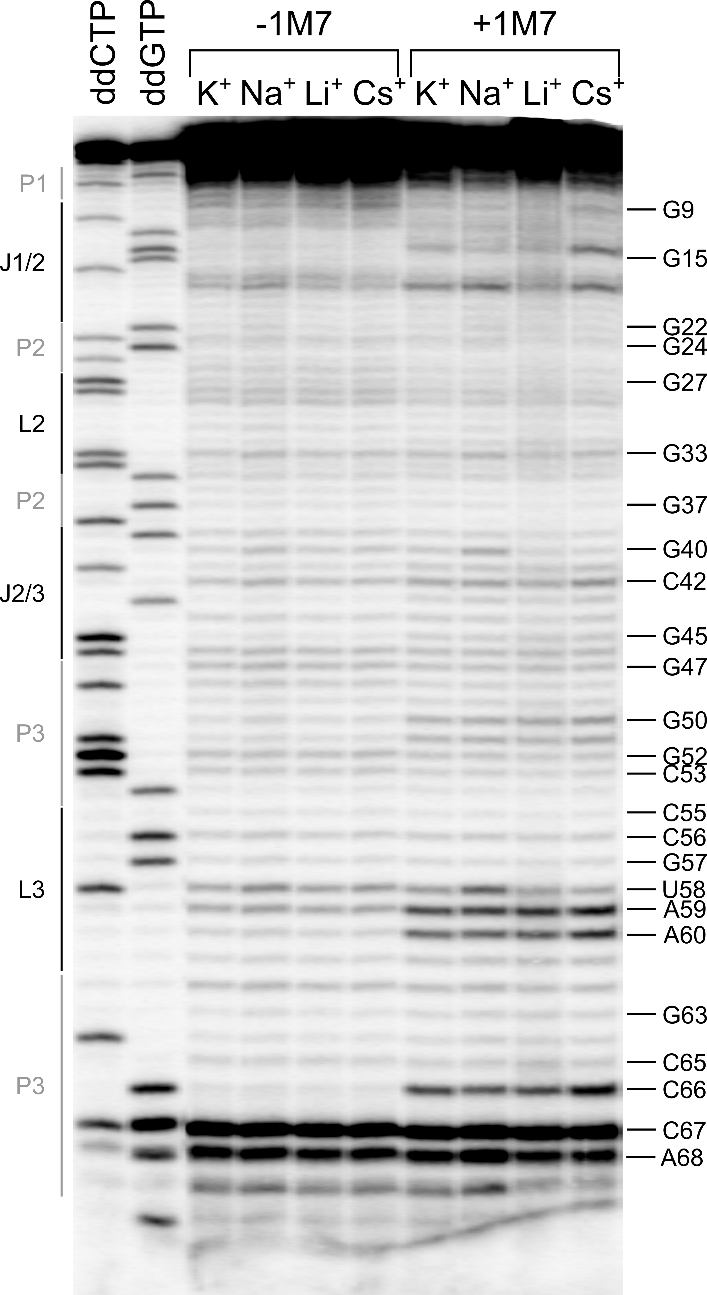

Supplementary Figure S7**.** DGR-1A footprinting in different monovalent cation buffers. The DGR-1A aptamer was reacted with 1M7 in different buffers containing either K^+^, Na^+^, Li^+^, or Cs^+^ to verify that the overall structure of the aptamer was not significantly perturbed by altering the monovalent cation identity. Across all buffers, the DGR-1A aptamer does not display significantly altered reactivity patterns.

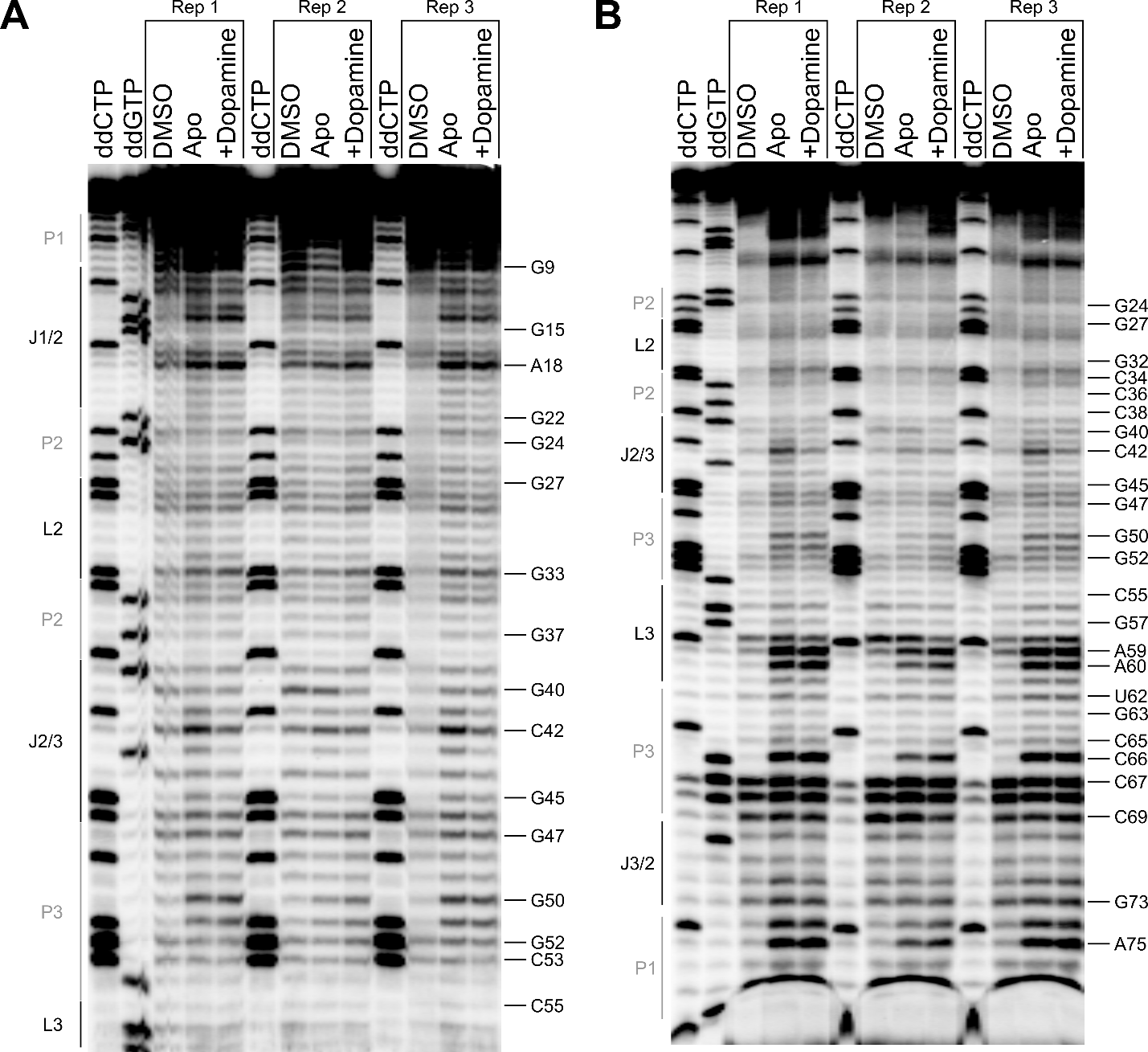

Supplementary Figure S8**.** Full DGR-1A footprinting gels with dopamine. The DGR-1A aptamer reacted with 1M7 in the presence or absence of 1 mM dopamine. Reactions were performed in triplicate. Samples were run on the gel for different amounts of time to resolve different regions of the RNA. The 5′ end of the RNA is resolved in (A) while the 3′ end is resolved in (B).

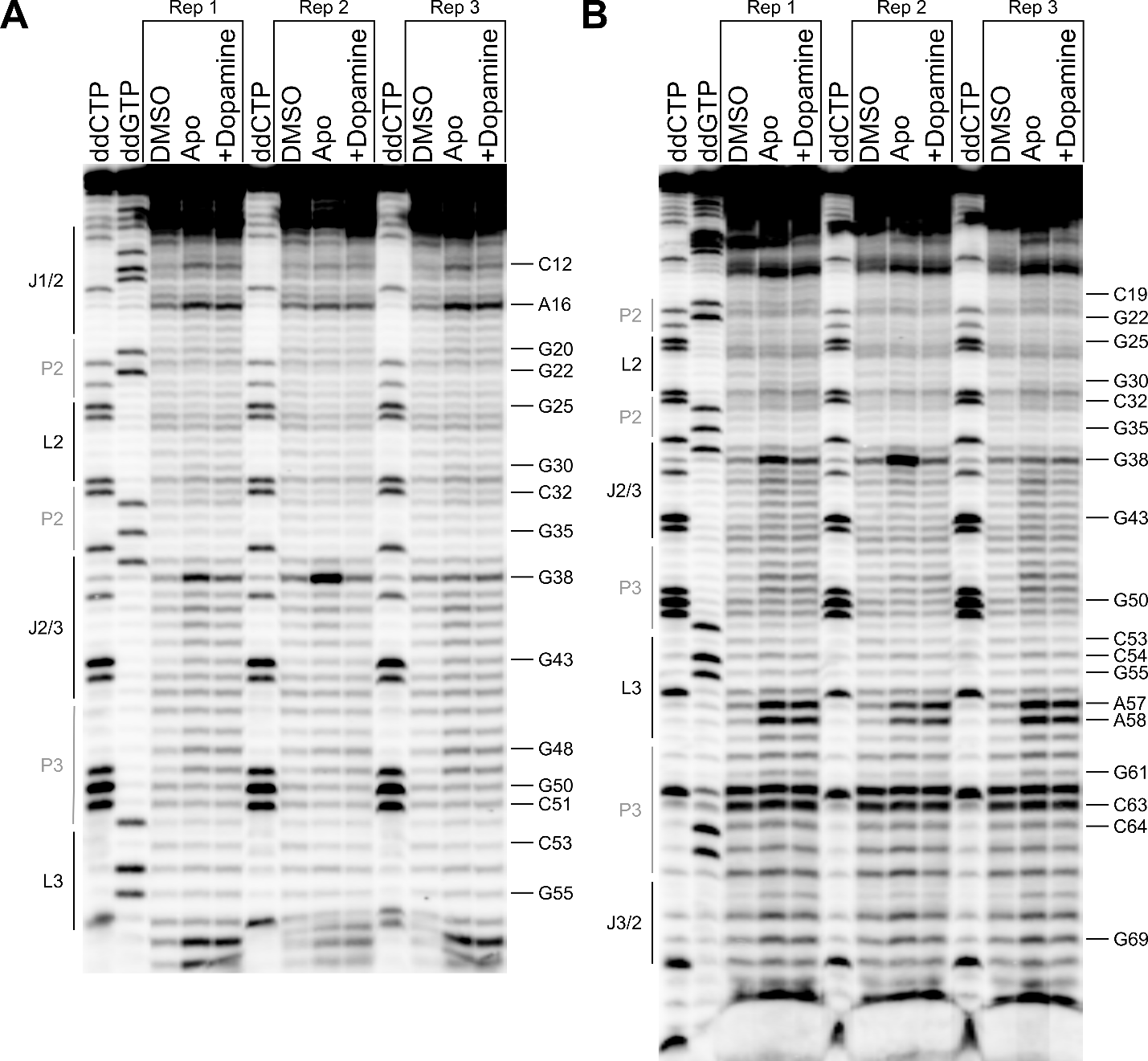

Supplementary Figure S9**.** Full footprinting gels of the DGR-1B aptamer. The DGR-1B aptamer reacted with 1M7 in the presence or absence of 1 mM dopamine. Reactions were performed in triplicate. Samples were run on the gel for different amounts of time to resolve different regions of the RNA. The 5′ end of the RNA is resolved in (A) while the 3′ end is resolved in (B).
